# The genetic organization of subcortical volumetric change is stable throughout the lifespan

**DOI:** 10.1101/2020.06.12.143834

**Authors:** Anders M Fjell, Håkon Grydeland, Yunpeng Wang, Inge Amlien, David Bartrés-Faz, Andreas M. Brandmaier, Sandra Düzel, Jeremy Elman, Carol Franz, Asta K. Håberg, Tim C. Kietzmann, Rogier A. Kievit, William S Kremen, Stine K Krogsrud, Simone Kühn, Ulman Lindenberger, Didac Macià, Athanasia M. Mowinckel, Lars Nyberg, Matthew S. Panizzon, Cristina Solé-Padullés, Øystein Sørensen, René Westerhausen, Kristine B Walhovd

## Abstract

While development and aging of the cerebral cortex show a similar topographic organization and are mainly governed by the same genes, it is unclear whether the same is true for subcortical structures, which follow fundamentally different ontogenetic and phylogenetic principles than the cerebral cortex. To test the hypothesis that genetically governed neurodevelopmental processes can be traced in subcortical structures throughout life, we analyzed a longitudinal magnetic resonance imaging dataset (n = 974, age 4-89 years), identifying five clusters of longitudinal change in development. With some exceptions, these clusters followed placement along the cranial axis in embryonic brain development, suggesting continuity in the pattern of change from prenatal stages. Developmental change patterns were conserved through the lifespan and predicted general cognitive function in an age-invariant manner. The results were replicated in longitudinal data from the Lifebrain consortium (n = 756, age 19-83 years). Genetic contributions to longitudinal brain changes were calculated from the Vietnam Era Twin Study of Aging (n = 331 male twins, age 51-60 years), revealing that distinct sets of genes tended to govern change for each developmental cluster. This finding was confirmed with single nucleotide polymorphisms and cross-sectional MRI data from the UK Biobank (n = 20,588, age 40-69), demonstrating significantly higher co-heritability among structures belonging to the same developmental clusters. Together, these results suggest that coordination of subcortical change adheres to fundamental principles of lifespan continuity, genetic organization and age-invariant relationships to cognitive function.

**Significance statement:** Here we show that subcortical change during childhood development is organized in clusters. These clusters tend to follow the main gradient of embryonic brain development, and are stable across life. This means that subcortical regions changing together in childhood also change together throughout the rest of life, in accordance with a lifespan perspective on brain development and aging. Twin and single nucleotide polymorphism-based heritability analyses in middle-aged and older adults showed that volume and volume change of regions within each developmental cluster tended to be governed by the same sets of genes. Thus, volumetric changes across subcortical regions are tightly organized, and the coordinated change can be described in a lifespan perspective according to ontogenetic and genetic influences.

## Introduction

Cortical development follows a topographic organization through childhood and adolescence (1-3). This topography is conserved through later development and aging (2, 3), closely following the genetic organization of the cortex (4). It is not known whether the same is true for subcortical structures. In contrast to the monotone thinning of the cerebral cortex (5, 6), lifespan trajectories of subcortical structures are more diverse and complex (7-11). This may be due to fundamental ontogenetic and phylogenetic differences between cortical and subcortical regions. The embryonic origin of the cortex is the pallium, while cerebellar and subcortical structures originate from the hindbrain, diencephalon or subpallium (12), and can be placed according to their position along the cranial vertical axis (see Table 2). Although the subcortex is evolutionary older than the cortex, it has a higher proportion of evolutionarily more recent genes, and a higher evolutionary rate, which is a basic measure of evolution at the molecular level (12). It has also been argued that genes expressed in the subcortex generally are more region-specific and tend to evolve more rapidly than genes expressed in cortical regions (12, 13). These mechanisms may be seen in human development and aging, with higher plasticity and potential for change in response to environmental impacts in phylogenetically older structures (14), especially the hippocampus (15, 16). This combination of plasticity and vulnerability could contribute to the larger diversity in the lifespan trajectories of subcortical structures (11). On the other hand, a hypothesis is that genetically governed neurodevelopmental processes can be traced in the brain later in life (17, 18), which has been shown for the comparably less plastic cortex (4). In light of the diverse age-trajectories and high plasticity of subcortical structures, it is not known whether patterns of subcortical maturation in childhood can be traced back to principles of embryonic development, how developmental organization sets constraints on subcortical aging, and the degree to which this organization of change is under common genetic control. The aim of the present study was to answer these unresolved issues about the organization of subcortical change across the lifespan. Specifically, we tested how subcortical developmental change clustered across different structures, how similar this organization was in development versus aging, and whether the patterns of change followed a common genetic anatomical architecture. We speculated that the developmental structures would tend to cluster according to embryonic principles, i.e. placement along the cranial vertical axis, that the pattern of change in aging would be similar to the pattern of change detected in childhood, and that structures changing together throughout the life would be governed by the same genes.

**Table 1.**
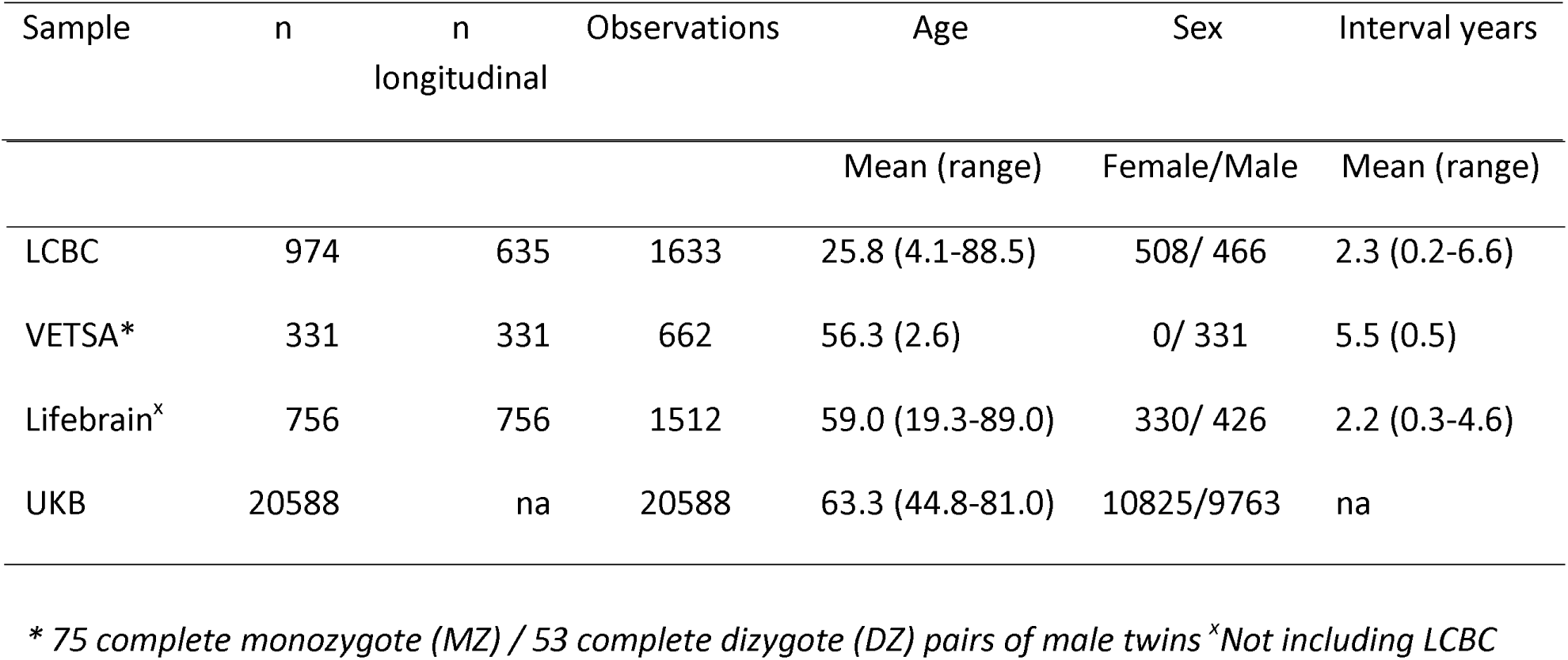
Sample overview.

**Table 2.** The embryonic origins of the clusters and placement along the cranial vertical axis

| Brain structure | Cluster | Embryonic development |  |  | Cranial vertical axis |
| --- | --- | --- | --- | --- | --- |
| 3rd ventricle | 1 | Prosencephalon (posterior) | Diencephalon |  |  |
| 4th ventricle | 1 | Rhombencephalon |  |  |  |
| Lat ventricle | 1 | Prosencephalon (anterior) | Telencephalon |  |  |
| Inf lateral ventricle | 1 |  |  |  |  |
| Brainstem (medulla oblongata) | 2 | Rhombencephalon | Myelencephalon |  | 0 |
| Cerebellum cortex | 2 | Rhombencephalon | Metencephalon |  | 1 |
| Cerebellum WM | 2 | Rhombencephalon | Metencephalon |  | 1 |
| Thalamus | 2 | Prosencephalon (posterior) | Diencephalon |  | 2 |
| Hippocampus | 2 | Prosencephalon (anterior) | Telencephalon (dorsal) | Pallium (medial) | 4 |
| Cortical WM | 2 | Prosencephalon (anterior) | Forebrain WM |  |  |
| Caudate | 4 | Prosencephalon (anterior) | Telencephalon (ventral) | Subpallium | 3 |
| Pallidum | 5 | Prosencephalon (anterior) | Telencephalon (ventral) | Subpallium | 3 |
| Putamen | 3 | Prosencephalon (anterior) | Telencephalon (ventral) | Subpallium | 3 |
| Accumbens | 3 | Prosencephalon (anterior) | Telencephalon (ventral) | Subpallium | 3 |
| Amygdala | 3 | Prosencephalon (anterior) | Telencephalon (dorsal) | Pallium (lateral) | 4 |
| Cortex | 3 | Prosencephalon (anterior) | Telencephalon (dorsal) | Pallium (dorsal) | 4 |

## Results

### Clusters of change in development adhere to embryonic development

First, we fitted trajectories of the volume of 16 brain regions to age, including the cortex, cerebellum, brain stem and the ventricles, with sex and intracranial volume (ICV) as covariates by Generalized Additive Mixed Models (GAMM)(19), using a single-center longitudinal dataset (Center for Lifespan Changes in Brain and Cognition [LCBC]), comprising 974 healthy participants from 4.1 to 88.5 years with a total of 1633 MRI examinations. The lifespan trajectories for the brain structures are presented in Figure 1 and statistics in SI. Next, the sample was divided into development and adulthood/ aging (development [< 20 years], n = 644, 1021 MRIs, follow-up interval = 1.7 years [1.0-3.2]; adulthood/ aging [≥ 20 years], n = 330, 612 MRIs, follow-up interval = 1.6 years [0.2-6.6]), see (4) for details (Sample descriptives are provided in Table 1). Annual symmetrized percent change (APC) was calculated for each participant for each brain region, averaged over hemispheres, using the formula APC = (Vol Tp2 – Vol Tp1) / (Vol Tp2 + Vol Tp1) × 100. These APCs were correlated between each pair of brain regions. The Louvain algorithm for detecting communities in networks was applied to derive clusters in the correlation matrix from the development sample, and the Mantel test run to compare the different matrices. Five clusters of coordinated developmental change were identified (Figure 2). Three large clusters consisted of the ventricles (Cluster 1), the brain stem, cerebellum white matter (WM) and cerebellum cortex, cortical WM, thalamus and hippocampus (Cluster 2) and cortex, putamen, amygdala and nucleus accumbens (Cluster 3). Caudate (Cluster 4) and pallidum (Cluster 5) were represented by separate clusters. An overview of the clusters and their embryonic developmental origins is given in Table 2. Although a one-to-one correspondence between embryonic development and clustering of change in childhood was not expected, there were clear tendencies to conservation of embryonic developmental principles in later childhood development. The regions of Cluster 2 mainly to emerged from rhombencephalon or the posterior prosencephalon, making up structures placed low on the cranial vertical axis, including brain stem (myelencephalon), cerebellum cortex and WM (metencephalon) and the thalamus (diencephalon). The exception to this was that the hippocampus and the cortical WM were also included in Cluster 2. The extensive connectivity between cerebellum and cerebrum and the existence of common factors of WM development may explain the latter finding. Clusters 3-5 comprised structures developed from subpallium/ventral telencephalon (caudate, pallidum, putamen) and pallium/dorsal encephalon (amygdala, cortex), also showing a certain consistency with the major principles from embryonic development and placement along the cranial vertical axis.

**Figure 1.**
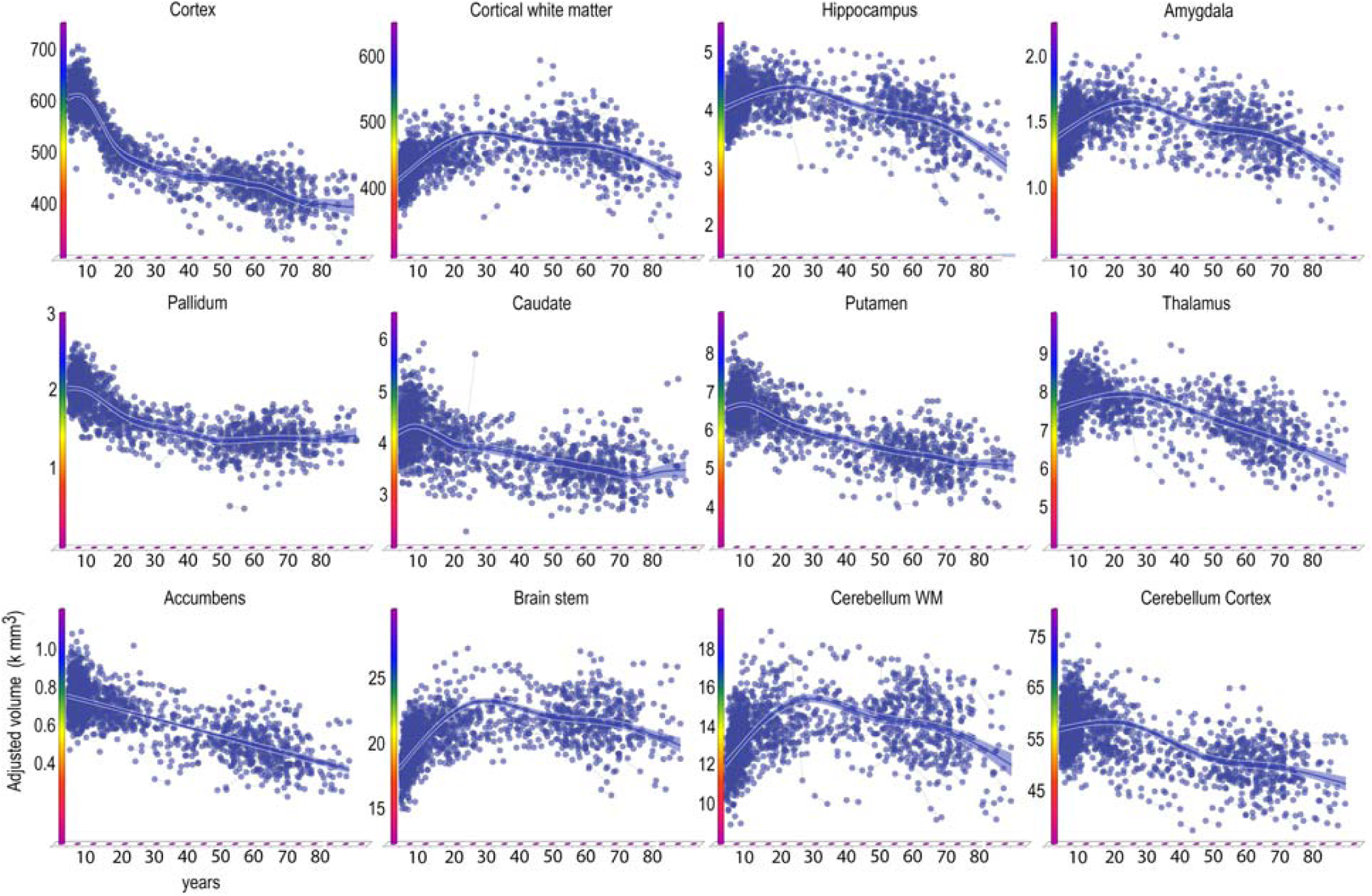
Lifespan trajectories of brain volumes. Age on the x-axis, volume in units of milliliters on the y-axis. The trajectories are fitted with GAMM, and the shaded areas represent 95% CI. Ventricular volumes not shown.

**Figure 2.**
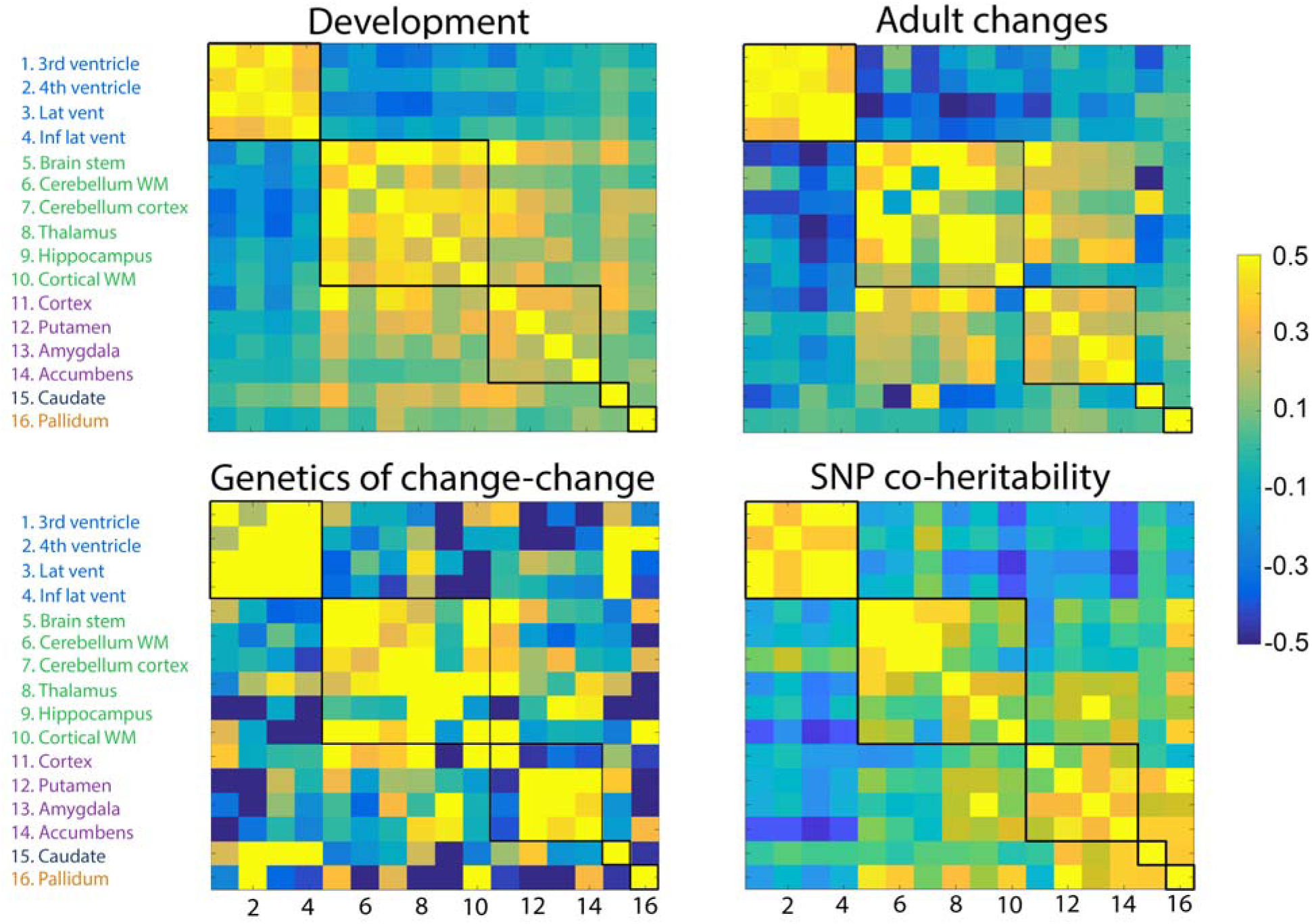
Volumetric change-change relationships. Top panels: Heat-maps representing pairwise correlations coefficients between volume change of the brain structures in development (left) and aging (right), i.e. change-change relationships. Bottom panels: The genetic change-change correlations (i.e. the genetic contribution to the relationships between change among any two structures) based on twin analysis (VETSA) are shown to the left, and the SNP co-heritability from the UKB cross-sectional data to the right. The five clusters were derived from the developmental sample.

We fitted the developmental trajectory of each cluster (Figure 3, statistics in SI). Cluster 1 increased linearly, although the rate of increase was modest. Cluster 2 showed a decreasing exponential function with volume increase levelling off after 15 years. Cluster 3 mimicked a cubic relationship, with a slight increase in volume until about 8 years, then steeper reductions, which were gradually smaller from 15 years. Cluster 4 (caudate) showed an inverted U-shaped trajectory with a sharp increase until about 9 years, and Cluster 5 (pallidum) a cubic trajectory with similarities to Cluster 3.

**Figure 3.**
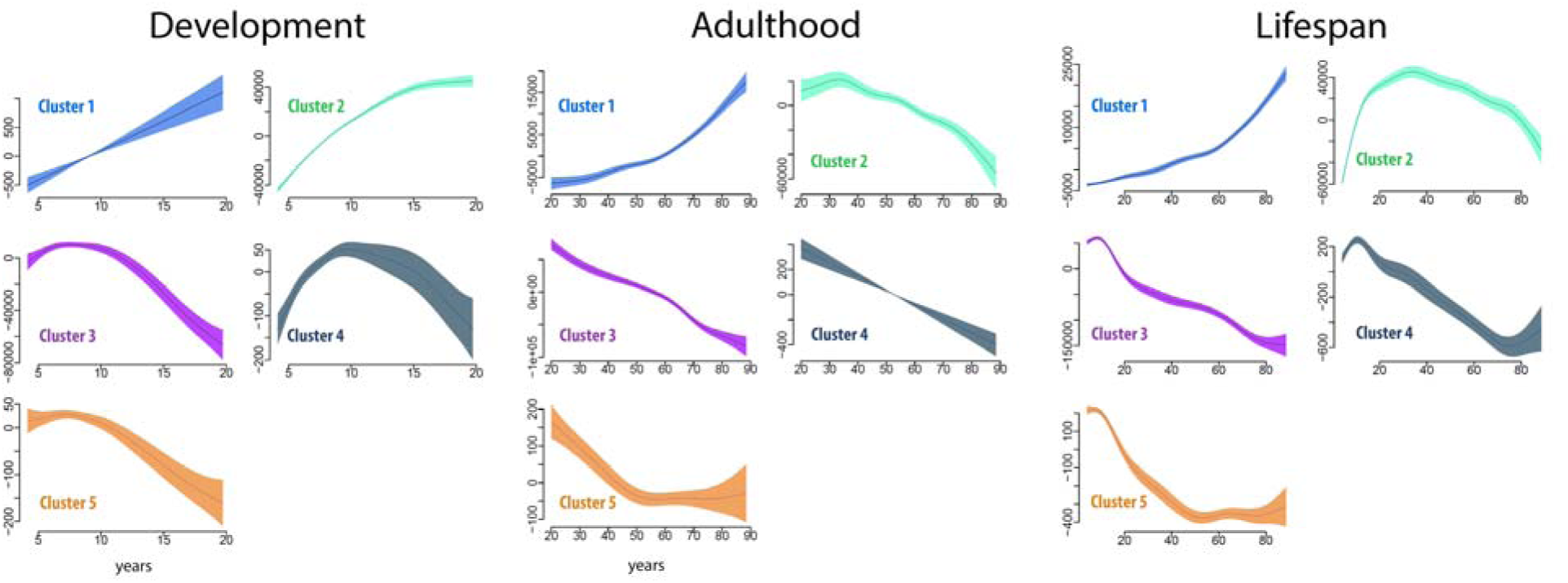
Cluster trajectories. Age-trajectories for each cluster, for development (left), adulthood (middle) and the full life-span (right). The trajectories are fitted with GAMM, and the shaded areas represent 95% CI. Note that the y-axes scales vary for easier viewing.

### Pattern of subcortical change in adulthood/ aging can be predicted from development

To test whether the topographical organization of change detected in development was conserved in adulthood and aging, the pairwise change-change correlations between regions were calculated for the adult and aging sample (Figure 2), and the Mantel test was run to compare the developmental and the adult/aging matrices. The change-change matrices were highly similar, r = .82 (p < .0001), demonstrating substantial overlap of clusters of change in development and aging. The results were replicated using longitudinal data from the Lifebrain consortium (total n = 756, 1512 MRIs, mean follow-up interval = 2.3 years, age 19-89 years, mean 59.8 years), yielding almost identical results (r = .78, p < .0001) (Figure 4). As an additional test, we ran two-sample Student’s t-tests, testing whether the mean correlation within the cluster identified in the developmental sample was larger than the mean correlation between the structures within the cluster and the structures outside the cluster (SI for details). The intra-cluster correlations were significantly higher than the extra-cluster correlations, both in the LCBC aging sample and the Lifebrain replication sample (Table 3). Finally, we ran the Mantel test comparing the patterns of developmental to adult and aging changes in 12 genetically defined cortical regions from the same LCBC participants (see (4, 20)). This yielded a correlation (r = .83) close to identical to what we observed for subcortex, suggesting that the organization of developmental subcortical change is conserved through life to the same degree as cortical change is.

**Table 3.** Within-vs. outside cluster correlations. Two-sample Student’s t-tests were run to test whether the mean correlation within the developmentally defined clusters was larger than the mean correlation between the variables in the cluster and the variables outside the cluster. Note that the clusters were defined in the developmental sample, which is independent from the other four samples.

| Dataset | Cluster 1<br>$P <$ | Cluster 2<br>$P \leq$ | Cluster 3<br>$P \leq$ |
| --- | --- | --- | --- |
| UKB (SNP co-heritability) | $1e^{-8}$ | 0.00002 | $6e^{-7}$ |
| VETSA (TWIN heritability) | $1e^{-8}$ | $3e^{-7}$ | 0.17 |
| Lifebrain (Aging) | $1e^{-8}$ | 0.0002 | 0.005 |
| LCBC (Aging) | $1e^{-8}$ | $1e^{-7}$ | $1e^{-7}$ |

**Figure 4.**
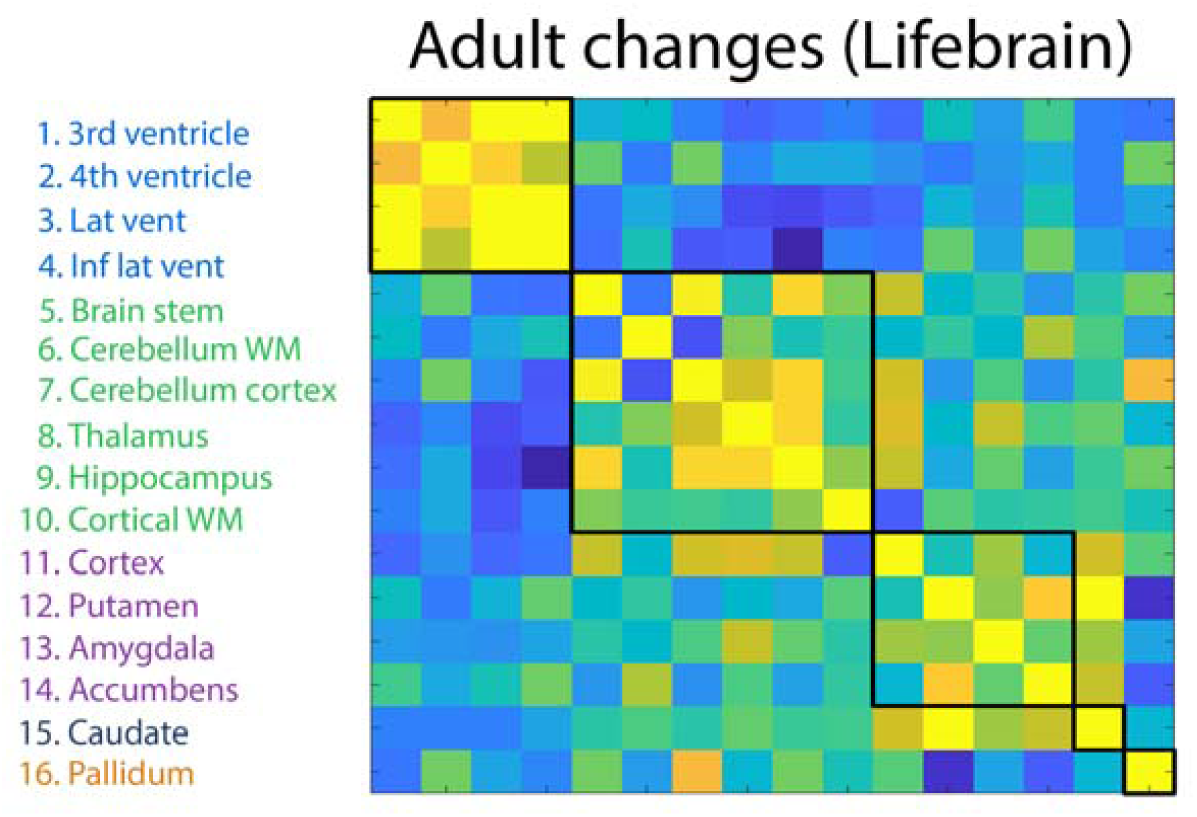
Lifebrain replication. Replication of the aging change-change pattern using Lifebrain data.

We calculated aging-trajectories for each of the clusters defined in the developmental sample and found that they differed qualitatively between clusters across the adult age-range. The trajectory for each cluster represented a continuation of the developmental trend seen in childhood and adolescence. Cluster 1 showed an exponential increase, cluster 2 an inverted U-shaped trajectory, cluster 3 and 4 almost linear reductions, while cluster 5 showed reductions until about 50 years and little or no change after that. For completeness, the age-trajectories of the clusters were also fitted across the full age-range from 4.1 to 88.5 years (Figure 3 lower right panel, numeric results in SI).

### Patterns of change adhere to principles of genetic organization

Using multivariate latent change score models, we estimated the differences in subcortical volumes from baseline to follow-up in the VETSA Twin data (n= 331, mean follow-up interval = 5.5 years) as well as pairwise genetic correlations of the slope factor (i.e., change) between all regions (see SI and (21) for details on the statistical twin model). This yielded an estimate of how much of the change-change relationship between any two brain structures is due to common genetic influence. The matrix of shared genetic influences on change between each pair of brain structures (Figure 2) was tested against the developmental and the adult/aging change-change correlation matrices. The Mantel test confirmed that the shared genetic influences on the change-change relationships were statistically more similar to the pattern of correlated changes during development (r = .39, p = .0014) and aging (r = .42, p = .0004) than expected by chance. Replication was again run using Lifebrain data, yielding r = .33 (p < .0035) between the matrix of shared genetic influences on change and the Lifebrain aging change-change correlation matrix. Mean intra-cluster genetic correlations were significantly higher than the extra-cluster genetic correlations for Cluster 1 and Cluster 2, but not for Cluster 3 (p = .17).

Finally, we calculated the pairwise single nucleotide polymorphism (SNP) genetic covariance between the volume of pairs of subcortical structures from the UK Biobank sample (n = 20,588, age 40-69 years), using age, sex and the first 10 components of the genetic ancestry factor as covariates. The pairwise genetic correlations are shown in SI. These analyses were based on cross-sectional data, not longitudinal change. Still, the SNP genetic correlation matrix was highly similar to the developmental change matrix, as demonstrated by the Mantel test (r = 0.71, p < .0001, see Figure 2) and significantly higher intra-than extra-cluster correlations (Table 3). This showed that the genetic organization of subcortical structures in middle-age can be predicted from the organization of change during brain development in childhood. For completeness, we also compared the SNP genetic matrix to the aging change matrix (r= 0.63, p < .0001) and the heritability change matrix from the VETSA sample (r = 0.36, p = .002), in both cases showing significantly higher similarity than expected by chance. The Lifebrain replication sample demonstrated excellent coherence with the UKB SNP co-heritability matrix (r = 0.61, p = .0005).

For completeness and to test relationship between principles of embryonic development and adult co-heritability, we ran hierarchical clustering on the UKB SNP co-heritability matrix to describe how shared genetic influences were distributed across structures (Figure 5 for details). This revealed a close to perfect match between the adult genetic clusters and embryonic origins. At the five cluster level, pallidum, putamen, nucleus accumbens and caudate clustered together, all originating from the subpallium (ventral telencephalon), which is the developmental origin of the basal ganglia. Further, amygdala, hippocampus and the cerebral cortex clustered together, all from their common known embryonic origin, the pallium (dorsal telencephalon). Brainstem, cerebellum WM and cerebellum cortex, all from the rhombencephalon, further clustered together. The ventricles formed a separate cluster, while thalamus and the cerebral WM constituted the last cluster. Although this analysis is auxiliary to the main change-analyses reported above, it demonstrates the same fundamental principle of close correspondence between the embryonic brain development and the brains genetic architecture decades later.

**Figure 5.**
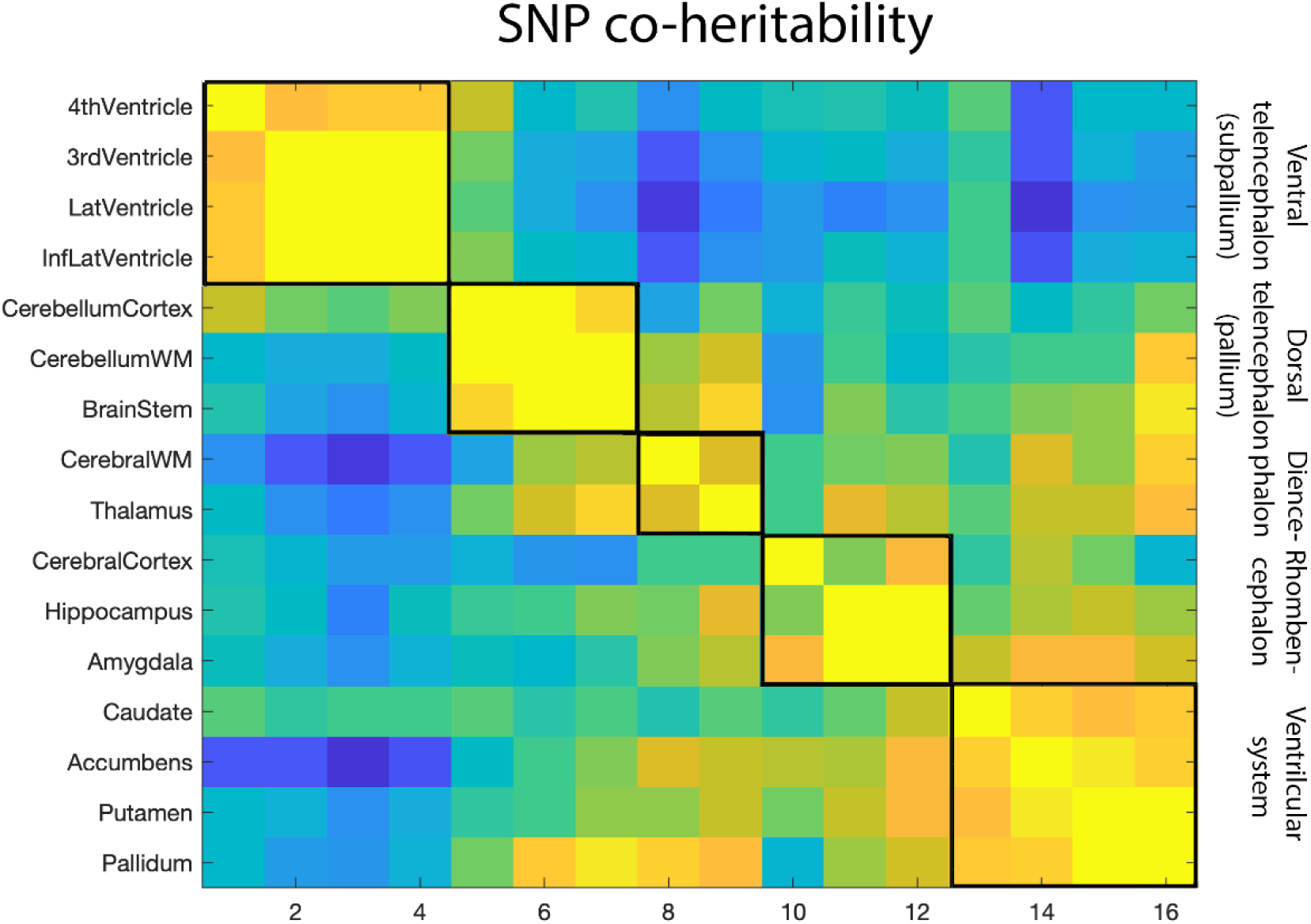
Correspondence between SNP heritability and embryonic brain development. Hierarchic clustering was used to test how shared genetic variance were distributed across structures (left column), and the clusters were compared to the main organization of embryonic brain development (right column). Hierarchic clustering was used to test how shared genetic variance were distributed across structures, and the clusters were compared to the main organization of embryonic brain development.

### Relationship to general cognitive function

Additional GAMMs were run to test the relationship between the development clusters and general cognitive function (GCA) as measured by the Wechsler’s Abbreviated Scale of Intelligence (22) in the full LCBC sample, using sex and age as covariates. Cluster 1 (t = 3.74, p = .00018), cluster 3 (t = 2.75, p = .006) and cluster 4 (t = 2.01, p = .0449) were significantly related to GCA. Low GCA score was associated with lower volumes. The relationship between GCA and clusters 1 and 3 survived including intracranial volume (ICV) as an additional covariate, while for Cluster 4 it was not significant (p = 0.071). Importantly, only for Cluster 1 was a significant interaction between GCA and age found (F = 4.59, p = .01), which was due to relatively less increase in ventricular volume in young adulthood in the low GCA group compared to the high GCA group. For the remaining clusters, the age-trajectories did not differ significantly as a function of GCA (all p’s ≥ .46), as can be seen in Figure 6.

**Figure 6.**
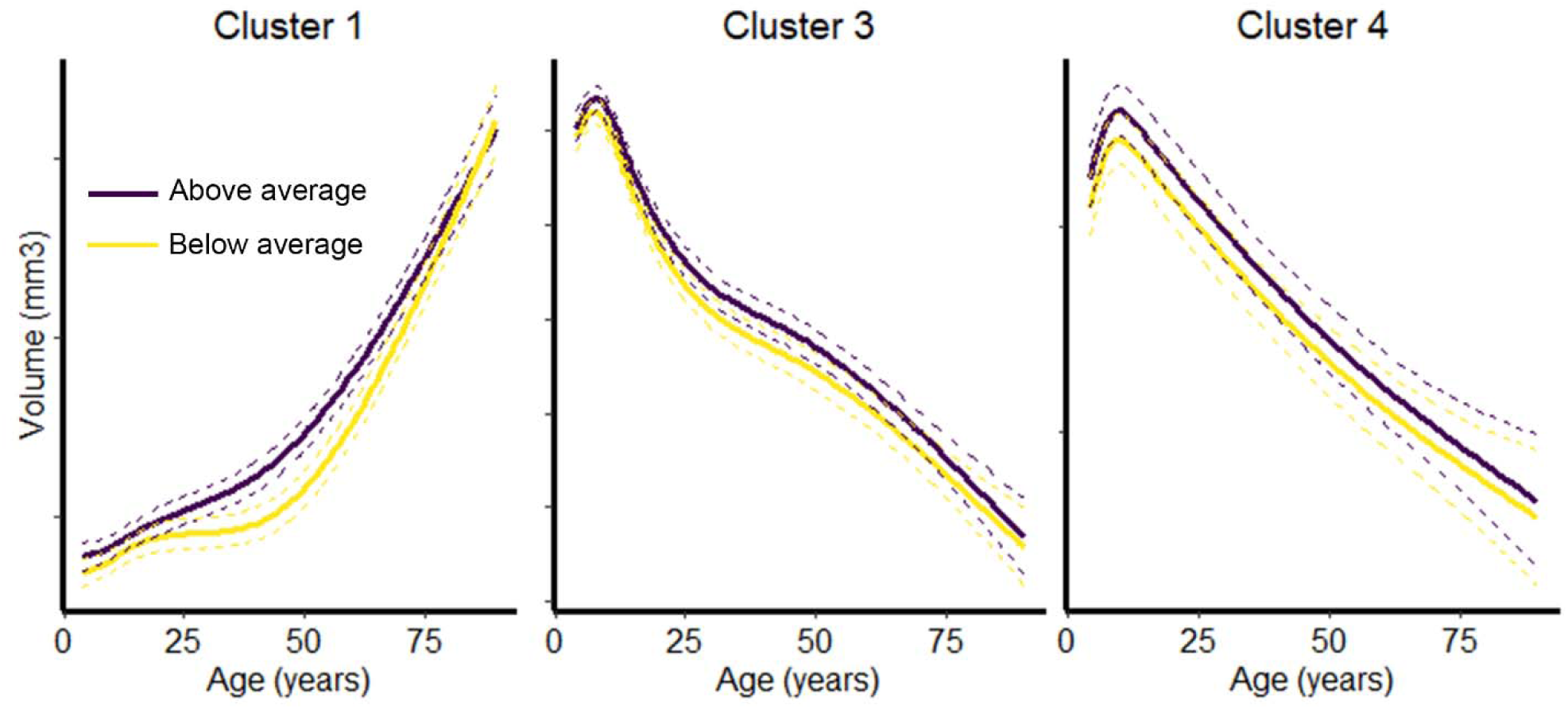
Age trajectories as a function of general cognitive function. Lifespan trajectories of the three clusters that were significantly related to general cognitive function. Trajectories are shown separately for participants above (purple lines) vs. below (yellow lines) the age-corrected median cognitive function. The trajectories are fitted with GAMM, and the dashed lines represent 95% CI.

## Discussion

The results demonstrate that subcortical structures develop in meaningful clusters that tend to follow the main cranial vertical axis from embryonic brain development. These clusters were conserved through life, and were governed by common sets of genes. Thus, although the lifespan trajectories of subcortical structures are much more divergent than the trajectories for cortical regions, clusters of change could be identified. The similarities to gradients of embryonic brain development suggest that there is continuity in the between-region pattern of change during childhood that may originate from much earlier stages in life. That the patterns of coordinated change in development also characterized aging-changes, and the stable, mostly age-invariant relationship to cognitive function, demonstrated consistency across life. Thus, similar to what has been shown for the cerebral cortex (4), regional brain aging can be predicted from developmental changes in childhood, which makes a strong case for the theory that early life sets the stage for aging (23-25). Analyses of a longitudinal sample of middle-aged twins further showed that each of the developmental clusters tended to be governed by the same genes. This conclusion was strengthened by the close similarity between the pattern of change in development and the SNP co-heritability for the same clusters in the UKB dataset. Thus, the results show that coordination of change across subcortical structures adheres to similar principles of lifespan continuity, genetic organization and relationships to cognitive function we previously have seen for cortical regions (3, 4, 23). This yields further support for the hypothesis that genetically governed neurodevelopmental processes can be traced in subcortical structures throughout life.

### Superordinate structures of change in childhood is linked to embryonic brain development

The topographical development of cortical thickness is well described, with monotone decreases and regional variability that seem to follow certain functional, anatomical and genetic principles of organization (1, 3, 4, 20, 26). The highly divergent lifespan trajectories of subcortical structures (7-9), the fundamental ontogenetic and phylogenetic differences between cortical and subcortical regions (12), and the differences in evolutionary rate and regional specificity of gene expression (12, 13), suggest that it is an open question whether the principles of cortical lifespan changes automatically can be applied to subcortical structures. On this background, the present results are intriguing. Five clusters of developmental change were found, which in large showed divergent developmental trajectories and showed resemblance to embryonic origins. For instance, the majority of Cluster 2 structures emerged from rhombencephalon or the posterior prosencephalon, which is the origin of structures placed low on the cranial vertical axis. This included the brain stem (myelencephalon), cerebellum cortex and cerebellum WM (metencephalon) and the thalamus (diencephalon). Clusters 3-5 comprise structures developed from subpallium/ ventral telencephalon (caudate, pallidum, putamen) and pallium/ dorsal encephalon (amygdala, cortex). Still, it is necessary to note departures from this principle, i.e. the inclusion of the hippocampus and the cortical WM in the same cluster. Hippocampus is part of the cerebral cortex, but develops from the medial pallium in contrast to the neocortex. Compared to many regions in the cerebral cortex, hippocampus has a more similar appearance across the range of mammal species (27). Further, cortical WM was placed in Cluster 2. Cortical WM originates from the anterior prosencephalon, but the major part of myelination occurs postnatally, so this compartment of the brain is not easily placed within the same embryonic developmental context. However, the WM label is anatomically gross, since there are regional differences in development (26). Thus, at a general level, change in the structures tended to cluster according to trends from embryonic development and placement along the cranial vertical axis, but with notable exceptions.

### Consistency in patterns of change across the lifespan and relationships to cognitive function

We tested the similarity between the developmental and the adult and aging change matrices. This showed substantial coherence in the pattern of change between development and the rest of the life, similar to previously observations for the cerebral cortex (4). This means that subcortical structures that develop together during childhood tend to change together in adulthood and aging. When the developmental clusters were mapped to the adult part of the sample, the change trajectories were clearly different across the rest of the age-span. Except clusters 3 and 4, which showed mostly linear negative trajectories, substantial qualitative differences in the shapes of the slopes were observed, suggesting that clusters identified in development continued to show independent trajectories of change through the rest of life. The lifespan trajectories showed expected shapes, with accelerated increases for the ventricular system (cluster 1) and an inverse U-shape for cluster 2, consisting of structures known to show this pattern, such as WM, hippocampus and the brain stem (7, 11). Clusters 3, 4 and 5 showed different variants of initial increase in childhood, followed by relatively linear decline through most of adulthood that leveled off at high age, with variations between clusters in terms of break points. Cluster 4 (caudate) and 5 (pallidum) consisted of only one structure each. Cluster 3, however, consisted of four structures (cortex, putamen, amygdala and nucleus accumbens), mimicking the known trajectory of cortical volume with more or less linear decline after the peak is reached in late childhood or early adolescence.

Clusters 1, 3 and to a lesser extent 4 were related to GCA in a mostly age-invariant manner, which implies that the relationship between cognitive function and subcortical volumes is established early in life. Relationships were expected, as meta-analyses have shown a robust - although modest - relationship between measures of intelligence and gross brain volume of r = .24 (28). Relationships between GCA and cluster 3 were expected, as cortical volume has been reported to correlate with GCA previously, including a study of the same sample using different measures of cortical structures (3, 23). The positive relationship between cluster 1, consisting of the ventricular system, and GCA was not expected, and warrants further examination in independent samples.

### Genetic influence on change

By estimating genetic contributions to the phenotypic change-change patterns, we were able to test whether change in regions that cluster together are influenced by the same genes. We found that the genetic correlation matrix was significantly more similar to the development matrix than what would have been expected by chance, and the genetic correlations within the development clusters were higher than the genetic correlations between the cluster regions and the outside regions. This means that regions that change together through life also tend to be governed by the same sets of genes. Thus, there seems to be a genetic basis for the consistent pattern of change in subcortical structures from development and through the rest of life. It must be noted that although significant heritability estimates for brain changes have been demonstrated in ENIGMA, evidence for the existence of genetic variants specific for brain change has been found only for cerebellar gray matter and the lateral ventricles (21). Rather, genetic influence on change rates and baseline volume overlapped for most structures. Thus, as an additional test we calculated SNP co-heritability based on UKB data. These data are cross-sectional, and thus do not yield information about change-change relationships per se. Still, we found strong evidence for regions within the development clusters being controlled by the same genes. A previous multi-sample GWAS also reported significant genetic correlations between some of the subcortical structures tested in the present study (18). The similarity of the developmental change matrix and the SNP co-heritability matrix obtained from middle-aged adults yields further support for the hypotheses that genetically governed neurodevelopmental processes can be traced in subcortical structures through life.

## Conclusion

Subcortical change during childhood development can be organized in clusters, which are stable through life and tend to follow gradients of embryonic brain development. Regions within each cluster are governed by similar sets of genes. Thus, the pattern of change in subcortical regions can be described in a lifespan perspective according to ontogenetic and genetic influences.

## Materials and Methods

### Samples

Multiple independent samples were used (Table 1). Details for all samples are found in SI.

#### (1) LCBC Lifespan sample

A total of 1633 valid scans from 974 healthy participants (508 females/ 466 males), 4.1 to 88.5 years of age (mean visit age 25.8, SD 24.1), were drawn from studies coordinated by the Research Group for Lifespan Changes in Brain and Cognition (LCBC), Department of Psychology, University of Oslo, Norway (4). For 635 participants, one follow-up scan was available while 24 of these had two follow-ups. Mean follow-up interval was 2.30 years (0.15-6.63 years, SD 1.19). Sample density was higher in childhood/ adolescence than adulthood, since we expected more rapid changes during that age period (1006 observations < 10 years, 378 observations ≥20 and < 60 years, and 249 observations 60-88.5 years). All participants’ scans were examined by a neuroradiologist, and deemed free of significant injuries or conditions. The studies were approved by the Regional Committee for Medical and Health Research Ethics South. Written informed consent was obtained from all participants older than 12 years of age and from a parent/guardian of volunteers under 16 years of age. Oral informed consent was obtained from all participants under 12 years of age.

#### (2) VETSA

331 male twins (150 MZ/106 DZ paired twins/75 unpaired) were randomly recruited from the Vietnam Era Twin Registry and had imaging data at two time points. Written informed consent was obtained from all participants. Average age at baseline was 56.3 (2.6) years and follow-up interval 5.5 (0.5) years (see (29, 30). Based on demographic and health characteristics, the sample is representative of US men in their age range (29, 31).

#### (3) The Lifebrain Consortium

756 participants with longitudinal MRI were included from the European Lifebrain project (1672 scans, baseline age 19-89 [mean = 59.8, SD = 16.4], mean follow-up interval 2.3 years, range 0.3-4.9, SD = 1.2) (http://www.lifebrain.uio.no/) (3), including major European brain studies: Berlin Study of Aging-II (BASE-II) (32, 33), the BETULA project (34), the Cambridge Centre for Ageing and Neuroscience study (Cam-CAN) (35), and University of Barcelona brain studies (36-38). Participants were screened to be cognitively healthy and in general not suffer from conditions known to affect brain function, such as dementia, major stroke, multiple sclerosis etc. Exact screening criteria were not identical across sub-samples.

#### (4) UK Biobank

20.588 participants with available MRIs and quality checked (QC) genetic information were included in the final analyses from UKB (40-69 years), see https://biobank.ndph.ox.ac.uk/. We received called genotypes for 488,377 subjects, of whom 21,916 had available MRIs pre-processed by FreeSurfer v6.0. We performed quality control of the genotype data at the participant level by removing participants failing genotyping QC (n=550), with abnormal heterozygocity values (n=969) or being relatives of other participants (n=515). In addition, we removed 481 participants suggested to be removed for genetic analysis by the UK Biobank team. At variant level, we removed SNPs having minor allele frequency less than 0.01 and end up with 65,4584 SNPs.

### Cognitive testing

General cognitive function was assessed by WASI (22) for participants aged 6.5-89 years of age, while scores for corresponding subtests (Vocabulary, Similarities, Block design and Matrices) from the Wechsler Preschool and Primary Scale of intelligence – III (WPPSI-III)(39) were used for the youngest participants (< 6.5 years) (see (23) and SI).

### MRI data acquisition and analysis

Imaging data for the LCBC sample were acquired using a 12-channel head coil on a 1.5-Tesla Siemens Avanto scanner (Siemens Medical Solutions, Erlangen, Germany) at Oslo University Hospital Rikshospitalet and St. Olav’s University Hospital in Trondheim. For all samples, subcortical volumes were obtained by use of FreeSurfer v6.0 (http://surfer.nmr.mgh.harvard.edu/) (40-42). For details regarding processing, scanner and sequence parameters for all samples, see SI.

### Genetic correlations

Genetic change correlations were obtained by latent change score analyses on the VETSA 1 and VETSA 2 subcortical data (see (21)). For the UKB SNP co-heritability analyses, the volume measures of the 16 sub-cortical structures were normalized to have zero mean and one standard deviation, separately, before estimating genetic correlations. We used the bivariate restricted maximum likelihood methods implemented in the program Genome-wide Complex Trait Analysis (GCTA(43)) to compute the genetic correlation for the volume measures for each pair of the 16 brain sub-cortical structures, including the first ten principal components, sex and age as covariates. The log likelihood test from GCTA testing whether a genetic correlation is zero was used to compute p values for estimated genetic correlations.

### Experimental Design and Statistical Analysis

GAMM implemented in R (www.r-project.org) using the package “mgcv” (19) was used to derive age-trajectories for all structures based on the 1633 LCBC MRIs. Annual symmetrized percent change (APC) in volume was correlated across structures in each sample separately (development and adult/ aging from LCBC and Lifebrain), and VETSA was used to calculate the genetic contribution to change-change relationships and SNP information for calculation of cross-sectional volume co-heritability in UKB. To identify clusters of correlations that could be compared across matrices, the community structure or modules in the matrices were obtained using the Louvain algorithm (44), part of the Brain Connectivity Toolbox (http://www.brain-connectivity-toolbox.net (45)). The optimal community structure is a subdivision of the network into non-overlapping groups of regions in a way that maximizes within-group connection strength, and minimizes between-group strength. The community structure may vary from run to run due to heuristics in the algorithm, so 10 000 iterations of the algorithm were run, and each region assigned to the module it was most often associated with (by taking the mode of the module assignment across iterations). To account for global brain changes, between-regional correlations were de-meaned before they were entered into the clustering analyses.

## Acknowledgements

The Lifebrain project is funded by the EU Horizon 2020 Grant: ‘Healthy minds 0–100 years: Optimising the use of European brain imaging cohorts (“Lifebrain”)’. Grant agreement number: 732592. Call: Societal challenges: Health, demographic change and well-being. In addition, the different sub-studies are supported by different sources:

LCBC: The European Research Council under grant agreements 283634, 725025 (to A.M.F.) and 313440 (to K.B.W.), as well as the Norwegian Research Council (to A.M.F., K.B.W.). Betula: a scholar grant from the Knut and Alice Wallenberg (KAW) foundation to L.N. University of Barcelona: Partially supported by a Spanish Ministry of Science, Innovation and Universities (MICIU/FEDER; RTI2018-095181-B-C21) to DB-F, which was also supported by an ICREA Academia 2019 grant award.; by the Walnuts and Healthy Aging study (http://www.clinicaltrials.gov; Grant NCT01634841) funded by the California Walnut Commission, Sacramento, California. BASE-II has been supported by the German Federal Ministry of Education and Research under grant numbers

16SV5537/16SV5837/16SV5538/16SV5536K/01UW0808/01UW0706/01GL1716A/01GL1716B, and S.K. has received support from the European Research Council under grant agreement 677804. Cam-CAN: Initial funding from the Biotechnology and Biological Sciences Research Council (BBSRC), followed by support from the Medical Research Council (MRC) Cognition & Brain Sciences Unit (CBU). VETSA is supported by U.S. National Institute on Aging grants R01s AG022381, and AG050595. Part of the research was conducted using the UK Biobank resource under application number 32048.

## Data availability

All code used in the analyses are available on request. The LCBC data supporting the results of the current study are available from the corresponding author on reasonable request, given appropriate ethical and data protection approvals. Requests for Lifebrain and VETSA data can be submitted to the relevant principal investigators of each study. Contact information can be obtained from the corresponding author. UK Biobank data requests can be submitted to www.ukbiobank.ac.uk.

