## Supplementary material for "The genetic organization of subcortical volumetric change is stable throughout the lifespan": SI

**for**

**1. Sample**

**2. Cognitive testing**

**3. Magnetic resonance imaging acquisition and preprocessing**

**4. GAMM fits lifespan**

**5. GAMM fits for the cluster trajectories**

**6. Heritability estimates for change-change in VETSA**

**7. UKB SNP co-heritability**

**8. Intra- vs. extra-cluster correlation tests**

**1. Sample**

Four samples were used in the present study.

*(1) LCBC Lifespan sample* Participants were drawn from three different studies coordinated by LCBC.

MoBa-Neurocog is a population based sample, with participants were recruited by the Norwegian Medical Birth Registry through the national Norwegian Mother and Child Cohort Study (1), see (2, 3). All participants in the cohort study living in the greater Oslo area or the greater Trondheim area were invited to participate in this study. Valid MRI scans were obtained from 472 participants (age 4.1 to 10.7 years at baseline, 231 girls and 241 boys), and since the project is conceptualized as a population-based community-sample study, all were included in the initial analyses. For 301 participants, valid scans at both baseline and follow up were acquired. 532 of the MoBa-Neurocog scans were performed in Oslo and 241 in Trondheim, on identical scanners with identical scanning parameters and sequences (see below).

Participants from NCD and CPLS were recruited through newspaper advertisements, and local schools and workplaces (n = 502). Detailed criteria for exclusion at baseline are described in (4, 5). The participants were screened using a standardized health interview prior to inclusion in the study. Participants with a history of self- or parent-reported neurological or psychiatric conditions, including clinically significant stroke, serious head injury, untreated hypertension, diabetes, and use of psychoactive drugs within the last two years, were excluded. Further, participants reporting worries concerning their cognitive status, including memory function, were excluded. All participants above 20 years of age scored <16 on Beck Depression Inventory (6) and participants above 40 years of age >26 on Mini Mental State Examination (7).

|  | Obs | n | Two Tps (n) | Three Tps (n) | Age^1^  (years) | Sex | Follow-up interval  (years) |
| --- | --- | --- | --- | --- | --- | --- | --- |
| Total sample | 1633 | 974 | 635 | 24 | 25.8 (4.1-88.5) | 508f/466m | 2.3 (0.2-6.6) |
| MoBa | 773 | 472 | 301 | 0 | 7.3 (4.1-12.0) | 231f/241m | 1.5 (1.0-2.2) |
| ND/ CPLS | 860 | 502 | 334 | 24 | 42.4 (8.2-88.5) | 277f/225m | 3.1 (0.2-6.6) |

^1^ Age at visit

Obs: Number of observations

n: Number of participants

Tp: Timepoint

MoBa: MoBa-Neurocog

**Table Sample characteristics for the LCBC sample**

*(2) Vietnam Era Study of Aging (VETSA)* The VETSA sample consisted of 331 male twins (150 MZ/106 DZ paired twins/75 unpaired), randomly recruited from the Vietnam Era Twin Registry, who had imaging data at two time points. Average age at baseline was 56.3 (2.6) years and follow-up interval 5.5 (0.5) (see (8, 9). The study was approved by the Institutional Review Boards at participating institutions, and all participants gave written informed consent. A total of 1237 male-male twins participated in wave one of this longitudinal study, and a subset underwent MRI. The MRI study began in year 3 of the primary study. Only 6% of those invited to participate in the MRI study declined to do so. Others were excluded for the following reasons: possible metal in the body (7%); claustrophobia (3%); inability to travel to the test site (5%); exclusion of co-twin (9%); other reasons (3%). Scanning problems resulted in a loss of data for 8% of the participants. The mean level of education was 13.9 years (±2.1). The narrow age range is part of a design focusing on longitudinal age-related change within a narrowly defined cohort. Based on demographic and health characteristics, the sample is representative of US men in their age range (9){Schoenborn, 2009 #36}. Variance in structural aspects of the brain in this sample is likely driven by several different factors, and previous studies have disentangled some of these, including testosterone (10, 11), cortisol (12) and cigarette smoking (13), as well as specific candidate genes, i.e. APOE (10) (14), in addition to general heritability (15).

*(3) The Lifebrain Consortium* The sample was derived from the European Lifebrain project (http://[www.lifebrain.uio.no/](http://www.lifebrain.uio.no/)) (16), including major European brain studies: Berlin Study of Aging-II (BASE-II) (17, 18), the BETULA project ((19), the Cambridge Centre for Ageing and Neuroscience study (Cam-CAN) (20), and University of Barcelona brain studies (21-23). Only participants with longitudinal MRIs available were included. LCBC data were not included in the Lifebrain analyses, as LCBC was used for exploration and Lifebrain for replication.

The studies included from Lifebrain were the following:

*Berlin Study of Aging-II* Participants were community-dwelling older adults recruited from the greater Berlin metropolitan area through advertisements in newspapers and public areas. Participants were recruited within the Berlin Aging Study II (BASE-II) (for cohort characteristics and additional details, see (17, 18)). None of the participants took medication that might affect memory function, and none had neurological disorders, psychiatric disorders, or a history of head injuries. All participants reported normal or corrected to normal vision, were right-handed, and scored over 27 on the Mini-Mental Status Examination. After completion the comprehensive cognitive examination of BASE-II, eligible participants were invited to take part in one MRI session within a time window of 2–4 weeks after cognitive testing. MRI and cognitive scores were obtained two times (baseline: 2012-2013; follow-up: 2015/2016), yielding a sample for the present analyses of 321 participants with longitudinal MRI (mean age at baseline = 61.9, SD = 16.7, range = 24.1-81.4, mean follow-up interval = 1.9 years, SD = 0.7, range = 0.6-3.1).

*The Betula longitudinal study on aging, memory and dementia* Population-based sampling was used for recruitment, detailed recruitment procedures are found in (19). The study is approved by the relevant ethical review board. Severe visual or auditory handicaps, intellectual or developmental disabilities, suspected dementia, having a mother tongue other than Swedish, MRI contraindications, neurological disorders, or visual/motor deficits that could interfere with fMRI data collection, MMSE <24, brain or head surgery, substantial brain anatomical deviations. Participation in the neuroimaging study was offered to all participants who had remained in the study and completed cognitive testing at the 5th Betula test wave in 2008-2009. A subset of 376 participants from the longitudinal Betula study underwent structural and functional MRI in 2009-2010 and 232 returned for a follow-up scan in 2013-2014. The parent samples from which the scanned participants were derived from were originally recruited to the study in 1988, 1993, and 2008 respectively. For the present analyses, longitudinal MRI data were available for 170 participants (mean age at baseline = 59.8, SD = 13.8, range = 25.5-80.8, mean follow-up interval = 4.0 years, SD = 0.2, range = 3.4-4.6).

*The Cambridge Centre for Ageing and Neuroscience (Cam-CAN) study* A population-based cohort of 3000 adults aged 18 was recruited to Stage 1 of the project, where they completed an interview including health and lifestyle questions, a core cognitive assessment, and a self-completed questionnaire of lifetime experiences and physical activity (20). Of those interviewed, ~700 participants aged 18-87 (100 per age decile) continued to Stage 2 where they undergo cognitive testing and provide measures of brain structure and function. A subset of ~250 adults returned for longitudinal follow-up data. General exclusion criteria: Term-time residents of colleges and universities, and participants whose Primary Care Physician feel are inappropriate to include.

Exclusion criteria for the MRI part of the study: Not cognitively normal (MMSE ≤ 24, memory defect, consent difficulties), communication difficulties (hearing problems [35db at 1000 Hz], insufficient English language, vision difficulties), medical problems by self-report of diagnosis (dementia diagnosis /Alzheimer’s Disease, Parkinson’s Disease, Motor Neuron disease, Multiple sclerosis, cancer, stroke, encephalitis, meningitis, epilepsy, head injury with serious results [coma, unconscious for >2 hours, skull fracture], recently diagnosed or uncontrolled high blood pressure, possible pregnancy, current psychiatric conditions [bipolar disorder, schizophrenia, psychosis]), mobility problems (restricted mobility which could prevent further participation, inability to walk 10 meters), substance abuse (past or current treatment for drug abuse, current drug usage), MRI/ MEG safety and comfort exclusions. The study is conducted in compliance with the Helsinki Declaration, and has been approved by the local ethics committee, Cambridgeshire 2 Research Ethics Committee (reference: 10/H0308/50). For the present analyses, longitudinal MRI data were available for 265 participants (mean age at baseline = 54.9, SD = 18.2, range = 19.3-89.0, mean follow-up interval = 1.4 years, SD = 0.7, range = 0.3-3.5).

*University of Barcelona brain studies* Healthy middle-aged/ older adults, all gave informed consent, in accordance with the Declaration of Helsinki (1964, last revision 2013). All study procedures were approved by the local Institutional Review Board). WAHA cohort: Eligible participants were recruited via mailing study brochures (LLU) or through the non-profit organization Institute of Aging (BCN), advertisements in the study centers, and word of mouth. Interested individuals attended an informational group meeting, completed a short medical questionnaire and signed the informed consent. CR/ iTBS cohorts: Healthy volunteers were recruited via the Institute of Aging, Barcelona. Individuals willing to participate were gathered in an informal meeting to tell them about the investigation, which included repetitive transcranial magnetic stimulation (TMS). GABA cohort: Participants were recruited from the Institute of Aging (Barcelona) and the University of Experience, an initiative by the University of Barcelona for students aged 55 and older, offering special one and two-year degrees. WAHA cohort: Participants were healthy elderly men and women with normal cognitive and visual function at the time of recruitment. Inclusion criteria were age between 63 and 79 years, apparently healthy, and equally willing to be in either of the two groups. Exclusion criteria included inability to undergo neuropsychological testing; morbid obesity (BMI ≥ 40 kg/m2); uncontrolled diabetes (HbA1c > 8%); uncontrolled hypertension (on-treatment blood pressure ≥ 150/100 mmHg); prior stroke, significant head trauma or brain surgery; relevant psychiatric illness; major depression; cognitive deterioration or dementia with a score < 24 on the Mini-Mental State Examination; other neurodegenerative disorders like Parkinson’s disease; advanced AMD or eye-related conditions precluding ophthalmological evaluation; prior chemotherapy; chronic illness with projected shortened lifespan; allergy to walnuts; customary use of fish oil and/or tree nuts (> 2 servings/week) and/or other relevant sources of ALA, such as flaxseed oil or soy lecithin. CR/iTBS cohort: Eligible participants had a normal cognitive profile with MMSE scores≥24 and performances not below 1.5SD according to normative scores (adjusted for age and education (Peña-Casanova et al., 2009)) on a neuropsychological evaluation that covered the major cognitive domains (including: Verbal memory: Rey auditory verbal learning test; visual memory: Rey-Osterrieth complex figure; Language: Benton naming test; semantic and phonetic fluencies; Frontal/Executive functions: direct and inverse digits, symbol digits modalities test, trail making test, Stroop test, London tower test; Visuospatial: line orientation, and visuoperceptive: Popplereuter’s embedded figures test). GABA cohort: None of the participants reported a diagnosis of a neurological or psychiatric disorder or any TMS contraindication (Rossi et al., 2009). Inclusion criteria for the older subjects included a normal cognitive profile with mini-mental state examination (MMSE; Folstein et al., 1975) scores of ≥24 and performance scores not more than 1.5 standard deviation (SD) below normative data (adjusted for age and years of education) on any of the administered neuropsychological tests (i.e., they did not fulfill the criteria for mild cognitive impairment (MCI); Petersen and Morris, 2005. The neuropsychological battery included (1) a screening test for dementia, using the MMSE, and an evaluation of: (2) premorbid cognition and intelligence quotient (IQ), using the vocabulary subtest of the Wechsler Adult Intelligence Scale-III (WAIS-III) and National Adult Reading Test (NART); (3) verbal memory, using the Free and Cued Selective Reminding Test (SRT); (4) executive functions, using the phonemic fluency task and Trail Making Test B (TMTB); (5) language, using the semantic fluency task and Boston Naming Test (BNT); and (6) speed of processing, using the Symbol Digit Modalities Test (SDMT). For the present analyses, longitudinal MRI data were available for 80 participants (mean age at baseline = 67.3 years, SD = 6.9, range = 36.8-78.1, mean follow-up interval = 3.7 years, SD = 0.9, range 1.6-4.9).

Key references: WAHA cohort: Rajaram S, Valls-Pedret C, Cofán M, Sabaté J, Serra-Mir M, Pérez-Heras AM, Arechiga A, Casaroli-Marano RP, Alforja S, Sala-Vila A, Doménech M, Roth I, Freitas-Simoes TM, Calvo C, López-Illamola A, Haddad E, Bitok E, Kazzi N, Huey L, Fan J, Ros E. The Walnuts and Healthy Aging Study (WAHA): Protocol for a Nutritional Intervention Trial with Walnuts on Brain Aging. Front Aging Neurosci. 2017 Jan 10;8:333.

CR/ iTBS cohorts: Vidal-Piñeiro D, Martin-Trias P, Arenaza-Urquijo EM, Sala-Llonch R, Clemente IC, Mena-Sánchez I, Bargalló N, Falcón C, Pascual-Leone Á, Bartrés-Faz D. Task-dependent activity and connectivity predict episodic memory network-based responses to brain stimulation in healthy aging. Brain Stimul. 2014 Mar-Apr;7(2):287-96.

GABA cohort: Abellaneda-Pérez K, Vaqué-Alcázar L, Vidal-Piñeiro D, Jannati A, Solana E, Bargalló N, Santarnecchi E, Pascual-Leone A, Bartrés-Faz D. Age-related differences in default-mode network connectivity in response to intermitent theta-burst stimulation and its relationships with maintained cognition and brain integrity in healthy aging. Neuroimage. 2018 Nov 22.

*(4)* *UK Biobank prospective epidemiological imaging study (all available data December 2019)* The UK Biobank data was released to LCBC by the application “Lifespan Changes in Brain and Cognition-identifying risk and protective factors” (project number 32048). See main text for details. UK Biobank is a prospective epidemiological resource gathering extensive questionnaires, physical and cognitive measures, and biological samples (including genotyping) in a cohort of 500,000 participants. An imaging extension to the existing UK Biobank study was funded in 2016 to scan 100,000 subjects from the existing cohort, aiming to complete by 2022 (24). Informed consent is obtained from all UK Biobank participants; ethical procedures are controlled by a dedicated Ethics and Guidance Council (http://www.ukbiobank.ac.uk/ethics) that has developed with UK Biobank an Ethics and Governance Framework (given in full at http://www.ukbiobank.ac.uk/wp-content/uploads/2011/05/EGF20082.pdf), with IRB approval also obtained from the North West Multi-center Research Ethics Committee.

**2. Cognitive testing**

General Cognitive Ability (GCA) was assessed by WASI (25) for participants aged 6.5-89 years of age, while scores for corresponding subtests (Vocabulary, Similarities, Block design and Matrices) from the Wechsler Preschool and Primary Scale of intelligence – III (WPPSI-III)(26) were used for the youngest participants (< 6.5 years) (see (27) and SI). All participants scored within normal IQ range (82-145), or normal range of scaled scores (mean of subtests, s = 6.67-17.33). The age-standardized GCA score was calculated by z-transforming each subtest according to age-group, and then calculating the mean of the z-scores. Since we had a very high sampling density in the younger age-ranges where we expected rapid cognitive development, z-transformations were performed separately for small age-ranges for the youngest participants, and then for gradually increasing age-ranges for older children and adults. Each age-group covered 0.5 (below 9 years of age), 1 (9-15 years), 2 (15-21), 5 (21-30 years) or 10 (30 years and above) years. If one sub-score was missing, (e.g. some subtests were rated by tester as non-valid for the youngest children) GCA would be computed based on the existing scores.

**3. Magnetic resonance imaging acquisition and preprocessing**

Details of the scanner and MRI acquisition parameters can be found in the table below.

| Sample | Scanner | Tesla | Sequence parameters |
| --- | --- | --- | --- |
| LCBC | Avanto Siemens | 1.5 | TR: 2400 ms, TE: 3.61 ms, TI: 1000 ms, flip angle: 8°, slice thickness: 1.2 mm, FoV: 240×240 m, 160 slices, iPat = 2 |
|  | Avanto Siemens | 1.5 | TR: 2400 ms, TE = 3.79 ms, TI = 1000 ms, flip angle = 8, slice thickness: 1.2 mm, FoV: 240 x 240 mm, 160 slices |
| Barcelona | Tim Trio Siemens | 3.0 | TR: 2300 ms, TE: 2.98, TI: 900 ms, slice thickness 1 mm, flip angle: 9°, FoV 256×256 mm, 240 slices |
| BASE-II | Tim Trio Siemens | 3.0 | TR: 2500 ms, TE: 4.77 ms, TI: 1100 ms, flip angle: 7°, slice thickness: 1.0 mm, FoV 256×256 mm, 176 slices |
| Betula | Discovery GE | 3.0 | TR: 8.19 ms, TE: 3.2 ms, TI: 450 ms, flip angle: 12°, slice thickness: 1 mm, FOV 250×250 mm, 180 slices |
| Cam-CAN | Tim Trio  Siemens | 3.0 | TR: 2250 ms, TE: 2.98 ms, TI: 900 ms, flip angle: 9°, slice thickness 1 mm, FOV 256×240 mm, 192 slices |
| UKB | Skyra  Siemens | 3.0 | TR: 2000 ms, TI: 880 ms, slice thickness: 1 mm, FoV: 208×256 mm, 256 slices, iPAT=2 |
| VETSA baseline | Siemens | 1.5 | TR=2730ms, TI=1000ms, TE=3.31ms, slice thickness=1.33mm, flip angle=7°, voxel size 1.3x1.0x1.3mm. Acquisition in Boston and San Diego. |
| VETSA follow-up (Boston) | Siemens Tim Trio | 3.0 | TE = 4.33 ms, TR = 2170 ms, TI = 1100 ms, flip angle = 7°, pixel bandwidth = 140, number of slices = 160, slice thickness = 1.2 mm. Acquisition in Boston. |
| VETSA follow-up (San Diego) | GE Discovery 750x | 3.0 | TE= 3.164  ms, TR = 8.084 ms, TI = 600 ms, flip angle = 8°, pixel bandwidth = 244.141, FOV = 24 cm, frequency = 256, phase = 192, number of slices = 172, slice thickness = 1.2 mm. Acquisition in San Diego. |

TR: Repetition time, TE: Echo time, TI: Inversion time, FoV: Field of View, iPat: in-plane acceleration

For LCBC data, the pulse sequences used for morphometric analysis were two repeated 3D T1-weighted magnetization prepared rapid gradient echo (MPRAGE). For the children between 4 and 9 years old in the MoBa-Neurocog sample, we used a parallel imaging technique (iPAT), acquiring multiple T1 scans within a short scan time, enabling us to discard scans with residual movement and average the scans with sufficient quality. Previous studies have shown that accelerated imaging does not introduce significant measurement bias in surface-based measures when using FreeSurfer for image analysis, compared with a standard MPRAGE protocol with otherwise identical voxel dimensions and sequence parameters (28), which is in accordance with our own analyses. The protocol also included a 25-slices coronal T2-weighted fluid-attenuated inversion recovery sequence (TR/TE =7000–9000/109 ms) to aid the neuroradiological examination.

*Preprocessing*

All MRI data were processed and analyzed with FreeSurfer (http://surfer.nmr.mgh.harvard.edu/) (29, 30). For LCBC and Lifebrain, to extract reliable volume estimates for each time point, images were automatically processed with the longitudinal stream (31) in FreeSurfer. Specifically an unbiased within-subject template space and image (32) is created using robust, inverse consistent registration (33). Several processing steps, such as skull stripping, Talairach transforms, atlas registration as well as spherical surface maps and parcellations are then initialized with common information from the within-subject template, significantly increasing reliability and statistical power (31). Volumes of each structure of interest were estimated based on well-established automated tools in FreeSurfer (34). FreeSurfer is an almost fully automated processing tool, and manual editing was not performed to avoid introducing errors. For children, the issue of movement is especially important, as it could potentially induce bias in the analyses (35). All children MRIs were manually rated for movement on a 1-4 scale, and only scans with ratings 1 and 2 (no visible or only very minor possible signs of movement) were included in the analyses, reducing the risk of movement affecting the results. Also, all reconstructed surfaces were inspected, and discarded if they did not pass internal quality control. This led to the exclusion of 46 participants from MoBa-Neurocog and 9 from ND, reducing the total sample to the reported 1633 scans. FreeSurfer 5.3 was used for the LCBC and VETSA analyses, while Lifebrain and UKB MRI data were processed with FreeSurfer 6.0. Because FreeSurfer is almost fully automated, to avoid introducing possible site-specific biases, gross quality control measures were imposed and no manual editing was done. Further details of the UKB imaging protocol (<http://biobank.ctsu.ox.ac.uk/crystal/refer.cgi?id=2367>) and structural image processing are provided on the UK biobank website (<http://biobank.ctsu.ox.ac.uk/crystal/refer.cgi?id=1977>).

**4. GAMM fits**

Numeric results for the GAMM fits for the lifespan LCBC data

|  | AIC | | BIC | | Effect of sex |
| --- | --- | --- | --- | --- | --- |
|  | Age | s(Age) | Age | s(Age) | p |
| Accumbens | 18324 | 18335 | 18357 | 18367 | 0.57 |
| Amygdala | 20360 | 20048 | 20392 | 20081 | 0.13 |
| Brainstem | 27514 | 26805 | 27546 | 26837 | 0.11 |
| Caudate | 22636 | 23558 | 22669 | 23591 | 0.75 |
| Cerebellum cortex | 30079 | 29946 | 30111 | 29979 | 0.94 |
| Cerebellum WM | 27430 | 27017 | 27463 | 27050 | 0.16 |
| Cortex | 37525 | 37895 | 37557 | 37927 | 0.72 |
| Cortical WM | 36847 | 36258 | 36879 | 36290 | 0.31 |
| Hippocampus | 22494 | 22235 | 22526 | 22268 | 0.22 |
| Pallidum | 20937 | 20741 | 20969 | 20774 | 0.17 |
| Thalamus | 23841 | 23669 | 23874 | 23701 | 0.06 |
| Total GM | 38130 | 37849 | 38163 | 37881 | 0.78 |
| Lateral ventricles | 30276 | 30147 | 30308 | 30174 | 0.33 |
| In flat vent | 21081 | 20931 | 21113 | 20964 | 0.23 |

**Table Generalized Additive Mixed Model fits LCBC Lifespan**

Generalized Additive Mixed Models (GAMM) were run with each neuroanatomical volume as dependent variable, and age, estimated total intracranial volume and sex as covariates. Separate models were run with a linear age (age) term or a slope function (s(Age)). Both Akaike Information Criterion (AIC) and Bayesian Information Criterion (BIC) were calculated to select among models and guard against over-fitting. In all cases yielded the slope function the lowest IC values, although for accumbens, the difference was not large. GM: Gray matter. WM: White matter.

**5. GAMM fits for the cluster trajectories**

|  | Development | | | Adulthood and aging | | | Lifespan | | |
| --- | --- | --- | --- | --- | --- | --- | --- | --- | --- |
|  | edf | F | p | edf | F | p | edf | F | p |
| Cluster 1 | 1.1 | 51.0 | .23e^-12^ | 6.0 | 67.1 | 2e^-16^ | 7.5 | 176.6 | 2e^-16^ |
| Cluster 2 | 6.5 | 363.5 | 2e^-16^ | 6.8 | 16.0 | 2e^-16^ | 8.7 | 214.3 | 2e^-16^ |
| Cluster 3 | 5.4 | 37.4 | 2e^-16^ | 6.7 | 85.5 | 2e^-16^ | 8.6 | 206.7 | 2e^-16^ |
| Cluster 4 | 5.7 | 18.1 | 2e^-16^ | 1.0 | 79.4 | 2e^-16^ | 8.3 | 45.9 | 2e^-16^ |
| Cluster 5 | 3.7 | 16.9 | 3.33e^-12^ | 3.8 | 15.8 | 1.23e^-11^ | 7.9 | 171.0 | 2e^-16^ |

**Table** Cluster age trajectories

Edf: effective degrees of freedom

**6. Heritability estimates for change-change in VETSA**

All subcortical volumes were adjusted for site and ICV. Left and right volumes at baseline and follow-up for each subject were included in a variant of the “latent change model” to characterize baseline subcortical volume and change in subcortical volume across the two assessment {McArdle, 2009 #65}, with the extension of modeling genetic and environmental effects on the phenotypes {Panizzon, 2015 #66}. The model allows for the estimation of the means and variances of the intercept and slope factors, the relative genetic (i.e., heritability) and environmental contributions to those variances, as well as the phenotypic, genetic, and environmental correlations between the latent factors. A genetic correlation matrix was generated by estimating genetic correlations of slope factors between all pairwise combinations of subcortical structures in bivariate latent change models.

**7. UKB SNP co-heritability**

| Trait1 | Trait2 | rg | se | P | sign p < .05 |
| --- | --- | --- | --- | --- | --- |
| Accumbens | Amygdala | 0.32 | 0.04 | 1.90E-13 | * |
| Accumbens | Brain_stem | 0.17 | 0.03 | 1.04E-06 | * |
| Accumbens | Caudate | 0.38 | 0.04 | < 2E-^16^ | * |
| Accumbens | Cerebellum_cortex | -0.02 | 0.04 | 0.26738 |  |
| Accumbens | Cerebellum_WM | 0.08 | 0.04 | 0.024322 | * |
| Accumbens | Cortex | 0.22 | 0.05 | 3.37E-06 | * |
| Accumbens | Cortical_WM | 0.27 | 0.04 | 5.65E-12 | * |
| Accumbens | Hippocampus | 0.20 | 0.04 | 1.65E-07 | * |
| Accumbens | Lat_ventrical | -0.43 | 0.04 | < 2E-^16^ | * |
| Accumbens | Pallidum | 0.39 | 0.04 | < 2E-^16^ | * |
| Accumbens | Putamen | 0.45 | 0.03 | < 2E-^16^ | * |
| Accumbens | Thalamus | 0.24 | 0.04 | 7.96E-09 | * |
| Accumbens | V3rd_ventrical | -0.33 | 0.04 | 2.74E-13 | * |
| Accumbens | V4th_ventrical | -0.33 | 0.04 | 1.85E-05 | * |
| Amygdala | Brain_stem | 0.04 | 0.03 | 0.11158 |  |
| Amygdala | Caudate | 0.24 | 0.04 | 3.57E-10 | * |
| Amygdala | Cerebellum_cortex | -0.01 | 0.04 | 0.40985 |  |
| Amygdala | Cerebellum_WM | -0.03 | 0.04 | 0.1978 |  |
| Amygdala | Cortex | 0.32 | 0.04 | 6.44E-11 | * |
| Amygdala | Cortical_WM | 0.17 | 0.04 | 9.28E-06 | * |
| Amygdala | Hippocampus | 0.51 | 0.03 | < 2E-^16^ | * |
| Amygdala | Lat_ventrical | -0.17 | 0.04 | 7.00E-05 | * |
| Amygdala | Pallidum | 0.26 | 0.04 | 5.35E-11 | * |
| Amygdala | Putamen | 0.32 | 0.03 | < 2E-^16^ | * |
| Amygdala | Thalamus | 0.23 | 0.04 | 8.10E-09 | * |
| Amygdala | V3rd_ventrical | -0.08 | 0.04 | 0.030717 | * |
| Amygdala | V4th_ventrical | -0.01 | 0.04 | 0.43368 |  |
| Brain_stem | Caudate | 0.10 | 0.03 | 0.000645 | * |
| Brain_stem | Cerebellum_cortex | 0.41 | 0.02 | < 2E-^16^ | * |
| Brain_stem | Cerebellum_WM | 0.75 | 0.02 | < 2E-^16^ | * |
| Brain_stem | Cortex | -0.18 | 0.04 | 2.08E-06 | * |
| Brain_stem | Cortical_WM | 0.23 | 0.03 | 6.70E-13 | * |
| Brain_stem | Hippocampus | 0.16 | 0.03 | 6.79E-07 | * |
| Brain_stem | Lat_ventrical | -0.17 | 0.04 | 3.11E-06 | * |
| Brain_stem | Pallidum | 0.45 | 0.03 | < 2E-^16^ | * |
| Brain_stem | Putamen | 0.19 | 0.03 | 1.30E-09 | * |
| Brain_stem | Thalamus | 0.40 | 0.03 | < 2E-^16^ | * |
| Brain_stem | V3rd_ventrical | -0.11 | 0.04 | 0.00149 | * |
| Brain_stem | V4th_ventrical | 0.02 | 0.03 | 0.24535 |  |
| Caudate | Cerebellum_cortex | 0.11 | 0.03 | 0.000432 | * |
| Caudate | Cerebellum_WM | 0.04 | 0.04 | 0.14967 |  |
| Caudate | Cortex | 0.06 | 0.04 | 0.094813 |  |
| Caudate | Cortical_WM | 0.03 | 0.03 | 0.21751 |  |
| Caudate | Hippocampus | 0.13 | 0.03 | 8.57E-05 | * |
| Caudate | Lat_ventrical | 0.08 | 0.04 | 0.020818 | * |
| Caudate | Pallidum | 0.37 | 0.03 | < 2E-^16^ | * |
| Caudate | Putamen | 0.33 | 0.03 | < 2E-^16^ | * |
| Caudate | Thalamus | 0.12 | 0.04 | 0.000533 | * |
| Caudate | V3rd_ventrical | 0.05 | 0.04 | 0.1188 |  |
| Caudate | V4th_ventrical | 0.11 | 0.03 | 0.000741 | * |
| Cerebellum_cortex | Cortex | -0.07 | 0.04 | 0.044272 | * |
| Cerebellum_cortex | Cortical_WM | -0.12 | 0.03 | 9.64E-05 | * |
| Cerebellum_cortex | Hippocampus | 0.07 | 0.03 | 0.012568 | * |
| Cerebellum_cortex | Lat_ventrical | 0.12 | 0.04 | 0.001542 | * |
| Cerebellum_cortex | Pallidum | 0.15 | 0.03 | 7.46E-06 | * |
| Cerebellum_cortex | Putamen | 0.06 | 0.03 | 0.02812 | * |
| Cerebellum_cortex | Thalamus | 0.14 | 0.03 | 2.21E-05 | * |
| Cerebellum_cortex | V3rd_ventrical | 0.15 | 0.04 | 4.19E-05 | * |
| Cerebellum_cortex | V4th_ventrical | 0.24 | 0.03 | 2.75E-13 | * |
| Cerebellum_WM | Cerebellum_cortex | 0.57 | 0.03 | < 2E-^16^ | * |
| Cerebellum_WM | Cortex | -0.17 | 0.05 | 0.000131 | * |
| Cerebellum_WM | Cortical_WM | 0.19 | 0.04 | 2.36E-07 | * |
| Cerebellum_WM | Hippocampus | 0.09 | 0.04 | 0.009874 | * |
| Cerebellum_WM | Lat_ventrical | -0.09 | 0.04 | 0.022321 | * |
| Cerebellum_WM | Pallidum | 0.36 | 0.03 | < 2E-^16^ | * |
| Cerebellum_WM | Putamen | 0.09 | 0.04 | 0.008247 | * |
| Cerebellum_WM | Thalamus | 0.26 | 0.04 | 4.16E-11 | * |
| Cerebellum_WM | V3rd_ventrical | -0.08 | 0.04 | 0.026293 | * |
| Cerebellum_WM | V4th_ventrical | -0.05 | 0.04 | 0.106 |  |
| Cortex | Cortical_WM | 0.08 | 0.04 | 0.036319 | * |
| Cortex | Hippocampus | 0.17 | 0.04 | 0.000125 | * |
| Cortex | Lat_ventrical | -0.14 | 0.05 | 0.002822 | * |
| Cortex | Pallidum | -0.05 | 0.04 | 0.14197 |  |
| Cortex | Putamen | 0.14 | 0.04 | 0.000406 | * |
| Cortex | Thalamus | 0.09 | 0.05 | 0.031157 | * |
| Cortex | V3rd_ventrical | -0.06 | 0.05 | 0.1278 |  |
| Cortex | V4th_ventrical | 0.02 | 0.04 | 0.36079 |  |
| Cortical_WM | Hippocampus | 0.15 | 0.03 | 2.59E-05 | * |
| Cortical_WM | Lat_ventrical | -0.41 | 0.04 | < 2E-^16^ | * |
| Cortical_WM | Pallidum | 0.39 | 0.03 | < 2E-^16^ | * |
| Cortical_WM | Putamen | 0.19 | 0.03 | 2.68E-08 | * |
| Cortical_WM | Thalamus | 0.28 | 0.03 | 3.28E-13 | * |
| Cortical_WM | V3rd_ventrical | -0.34 | 0.04 | 2.22E-16 | * |
| Cortical_WM | V4th_ventrical | -0.18 | 0.03 | 2.57E-07 | * |
| Hippocampus | Lat_ventrical | -0.23 | 0.04 | 5.61E-08 | * |
| Hippocampus | Pallidum | 0.19 | 0.03 | 7.86E-08 | * |
| Hippocampus | Putamen | 0.24 | 0.03 | 1.43E-12 | * |
| Hippocampus | Thalamus | 0.29 | 0.03 | 1.91E-14 | * |
| Hippocampus | V3rd_ventrical | -0.03 | 0.04 | 0.25557 |  |
| Hippocampus | V4th_ventrical | 0.02 | 0.04 | 0.24155 |  |
| Inf_vent | Accumbens | -0.36 | 0.05 | 1.29E-09 | * |
| Inf_vent | Amygdala | -0.08 | 0.06 | 0.090285 |  |
| Inf_vent | Brain_stem | -0.05 | 0.05 | 0.13087 |  |
| Inf_vent | Caudate | 0.09 | 0.05 | 0.046039 | * |
| Inf_vent | Cerebellum_cortex | 0.16 | 0.05 | 0.000714 | * |
| Inf_vent | Cerebellum_WM | -0.03 | 0.06 | 0.29009 |  |
| Inf_vent | Cortex | -0.15 | 0.06 | 0.012497 | * |
| Inf_vent | Cortical_WM | -0.33 | 0.05 | 2.77E-10 | * |
| Inf_vent | Hippocampus | -0.01 | 0.05 | 0.44615 |  |
| Inf_vent | Lat_ventrical | 0.73 | 0.03 | < 2E-^16^ | * |
| Inf_vent | Pallidum | -0.08 | 0.05 | 0.081325 |  |
| Inf_vent | Putamen | -0.09 | 0.05 | 0.036949 | * |
| Inf_vent | Thalamus | -0.19 | 0.05 | 0.000523 | * |
| Inf_vent | V3rd_ventrical | 0.57 | 0.04 | < 2E-^16^ | * |
| Inf_vent | V4th_ventrical | 0.37 | 0.05 | 5.02E-12 | * |
| Lat_ventrical | Pallidum | -0.17 | 0.04 | 3.89E-05 | * |
| Lat_ventrical | Putamen | -0.18 | 0.04 | 2.91E-06 | * |
| Lat_ventrical | Thalamus | -0.24 | 0.04 | 2.72E-08 | * |
| Lat_ventrical | V3rd_ventrical | 0.61 | 0.03 | < 2E-^16^ | * |
| Lat_ventrical | V4th_ventrical | 0.36 | 0.04 | < 2E-^16^ | * |
| Pallidum | Putamen | 0.61 | 0.02 | < 2E-^16^ | * |
| Pallidum | Thalamus | 0.34 | 0.03 | < 2E-^16^ | * |
| Pallidum | V3rd_ventrical | -0.14 | 0.04 | 0.000277 | * |
| Pallidum | V4th_ventrical | -0.04 | 0.04 | 0.10906 |  |
| Putamen | Thalamus | 0.24 | 0.03 | 1.26E-11 | * |
| Putamen | V3rd_ventrical | -0.07 | 0.04 | 0.037875 | * |
| Putamen | V4th_ventrical | -0.04 | 0.03 | 0.1411 |  |
| Thalamus | V3rd_ventrical | -0.18 | 0.04 | 1.23E-05 | * |
| Thalamus | V4th_ventrical | -0.02 | 0.04 | 0.30314 |  |
| V3rd_ventrical | V4th_ventrical | 0.33 | 0.04 | 8.33E-16 | * |

**SNP based heritability estimates** Pairwise co-heritability between brain structures derived from 20.588 participants from UKB. Age, sex, ICV and genetic ancestry (the first 10 components) were used as covariates. * p < .05 (uncorrected)

**8. Intra- vs. extra-cluster correlation tests**

Cluster Tests

library(readxl)
library(tidyverse)

We define the clusters in a list.

```
clusters <- list(
  Cluster1 = c("Bilatv3rdVentricle", "Bilatv4thVentricle", "BilatLatVentricle",
               "BilatInfLatVentricle"),
  Cluster2 = c("BilatBrainStem", "BilatCerebellumWM", "BilatCerebellumCortex",
               "BilatThalamus", "BilatHippocampus", "BilatCerebralWM"),
  Cluster3 = c("BilatCerebralCortex", "BilatPutamen", "BilatAmygdala",
               "BilatAccumbens"),
  Cluster4 = "BilatCaudate",
  Cluster5 = "BilatPallidum"
)
```

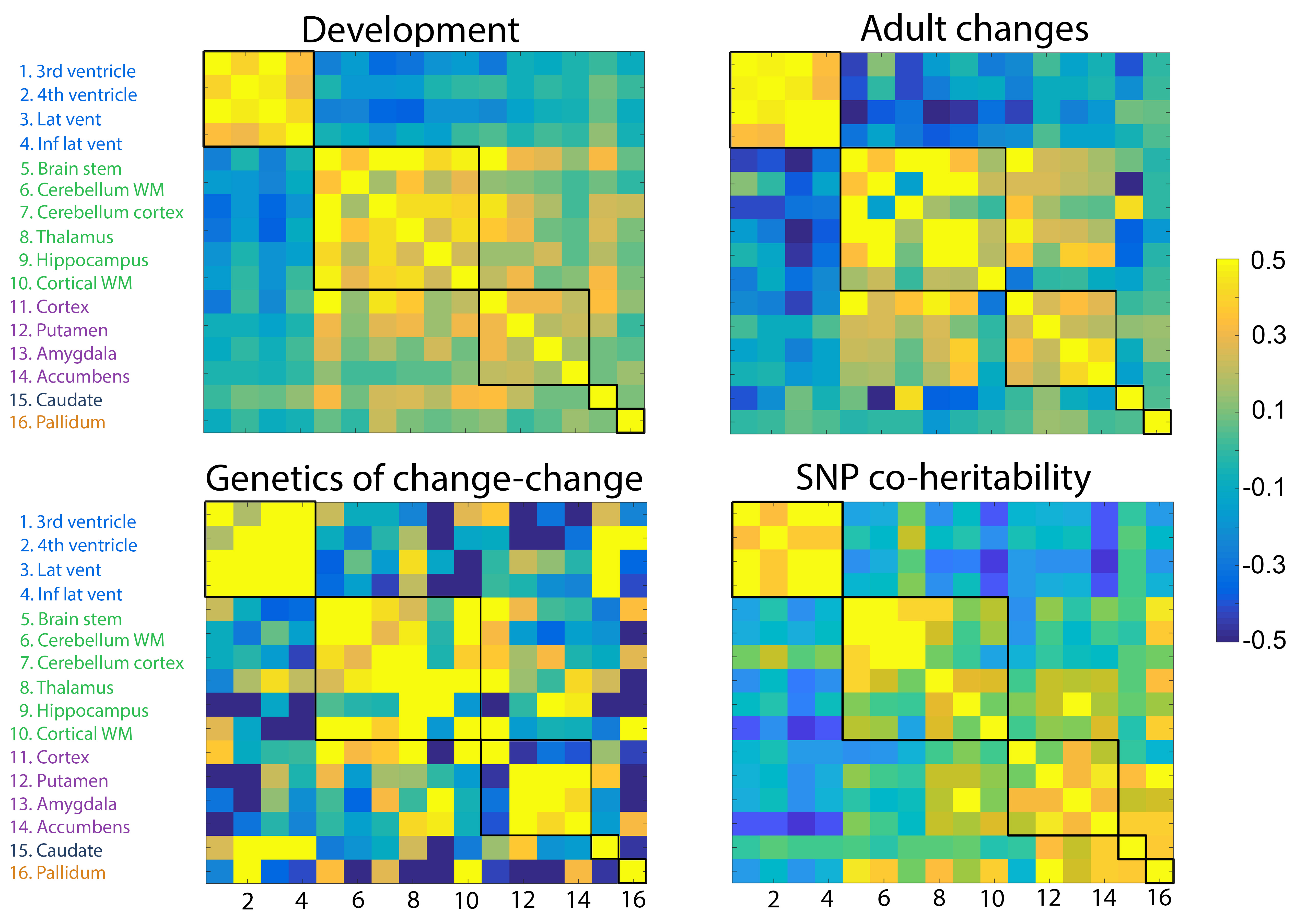

Function for testing correlations.

```
test_correlations <- function(dataset){
  imap(clusters, function(cl, nm){
    if(length(cl) > 1){
      correlations <- dataset %>%
        filter(Structure %in% cl) %>%   ##  Subset  to the right  rows
        pivot_longer(cols = -Structure, names_to = "Structure2",
                     values_to = "correlation") %>%  #  Reshape data
        filter(Structure != Structure2) %>% # Remove diagonal
        mutate( # Define whether it is  within or outside cluster
          within_cluster = Structure2 %in% cl
        ) %>%
        select(within_cluster, correlation) %>%
        group_by(within_cluster) %>%
        summarise(correlations = list(correlation))

      within_correlations <- correlations %>%
        filter(within_cluster) %>%
        pull(correlations) %>%
        unlist()

      between_correlations <- correlations %>%
        filter(!within_cluster) %>%
        pull(correlations) %>%
        unlist()

      cat("Testing in ", nm, "\n")

      test <- t.test(
        x = within_correlations,
        y = between_correlations,
        alternative = "greater"
      )

      print(test)
      cat("\n\n")

      return(test$p.value)
    } else {
      cat(nm, "has only a single member. No test performed.\n\n")
      return(NA_real_)
    }

  })
}
```

### UK Biobank

Read in the data.

```
ukb <- read_excel("UKB_GeneticCorr_Change_symmetric.xlsx") %>%
  rename(Structure = "...1") %>%
  rename_at(vars(-Structure), ~ paste0("Bilat", .)) %>%
  mutate(Structure = paste0("Bilat", Structure))
## New names:
## * `` -> ...1
```

Perform two-group t-tests in each cluster, testing if the correlation is larger in within than between. Information about the test is printed below.

```
ukb_tests <- test_correlations(ukb)
## Testing in  Cluster1
##
##  Welch Two Sample t-test
##
## data:  within_correlations and between_correlations
## t = 11.809, df = 16.669, p-value = 8.082e-10
## alternative hypothesis: true difference in means is greater than 0
## 95 percent confidence interval:
##  0.505255      Inf
## sample estimates:
##   mean of x   mean of y
##  0.49603817 -0.09661892
##
##
##
## Testing in  Cluster2
##
##  Welch Two Sample t-test
##
## data:  within_correlations and between_correlations
## t = 4.5152, df = 54.355, p-value = 1.72e-05
## alternative hypothesis: true difference in means is greater than 0
## 95 percent confidence interval:
##  0.129033      Inf
## sample estimates:
##  mean of x  mean of y
## 0.25743213 0.05241987
##
##
##
## Testing in  Cluster3
##
##  Welch Two Sample t-test
##
## data:  within_correlations and between_correlations
## t = 5.6833, df = 40.223, p-value = 6.471e-07
## alternative hypothesis: true difference in means is greater than 0
## 95 percent confidence interval:
##  0.1721583       Inf
## sample estimates:
##  mean of x  mean of y
## 0.29587617 0.05124969
##
##
##
## Cluster4 has only a single member. No test performed.
##
## Cluster5 has only a single member. No test performed.
```

### VETSA / Genetics of change-change

Read in the data.

```
# VETSA
vetsa <- read_delim("Subcort_GeneticCorr_Change_ed.csv", delim = ";",
                    locale = locale(decimal_mark = ",")) %>%
  rename(Structure = X1)
## Warning: Missing column names filled in: 'X1' [1]
## Parsed with column specification:
## cols(
##   X1 = col_character(),
##   BilatThalamus = col_double(),
##   BilatCaudate = col_double(),
##   BilatPutamen = col_double(),
##   BilatPallidum = col_double(),
##   BilatHippocampus = col_double(),
##   BilatAmygdala = col_double(),
##   BilatAccumbens = col_double(),
##   BilatInfLatVentricle = col_double(),
##   BilatLatVentricle = col_double(),
##   BilatCerebralWM = col_double(),
##   BilatCerebralCortex = col_double(),
##   BilatCerebellumCortex = col_double(),
##   BilatCerebellumWM = col_double(),
##   Bilatv3rdVentricle = col_double(),
##   Bilatv4thVentricle = col_double(),
##   BilatBrainStem = col_double()
## )
```

Perform two-group t-tests in each cluster, testing if the correlation is larger in within than between. Information about the test is printed below.

```
vetsa_tests <- test_correlations(vetsa)
## Testing in  Cluster1
##
##  Welch Two Sample t-test
##
## data:  within_correlations and between_correlations
## t = 8.2789, df = 36.711, p-value = 3.21e-10
## alternative hypothesis: true difference in means is greater than 0
## 95 percent confidence interval:
##  0.5412866       Inf
## sample estimates:
##  mean of x  mean of y
##  0.5668262 -0.1130322
##
##
##
## Testing in  Cluster2
##
##  Welch Two Sample t-test
##
## data:  within_correlations and between_correlations
## t = 5.5531, df = 67.32, p-value = 2.578e-07
## alternative hypothesis: true difference in means is greater than 0
## 95 percent confidence interval:
##  0.3130808       Inf
## sample estimates:
##   mean of x   mean of y
##  0.39946907 -0.04800671
##
##
##
## Testing in  Cluster3
##
##  Welch Two Sample t-test
##
## data:  within_correlations and between_correlations
## t = 0.98457, df = 13.426, p-value = 0.1711
## alternative hypothesis: true difference in means is greater than 0
## 95 percent confidence interval:
##  -0.1400012        Inf
## sample estimates:
##   mean of x   mean of y
##  0.15888474 -0.01736666
##
##
##
## Cluster4 has only a single member. No test performed.
##
## Cluster5 has only a single member. No test performed.
```

### Lifebrain

Translation of names for Lifebrain.

```
translate_lifebrain <- function(old_name){
  lifebrain_translation <- list(
    Accumbensarea_APC = "BilatAccumbens",
    Amygdala_APC = "BilatAmygdala",
    BrainStem_APC = "BilatBrainStem",
    Caudate_APC = "BilatCaudate",
    CerebellumCortex_APC = "BilatCerebellumCortex",
    CerebellumWhiteMatter_APC = "BilatCerebellumWM",
    CortexVol_APC = "BilatCerebralCortex",
    CerebralWhiteMatterVol_APC = "BilatCerebralWM",
    Hippocampus_APC = "BilatHippocampus",
    InfLatVent_APC = "BilatInfLatVentricle",
    LateralVentricle_APC = "BilatLatVentricle",
    Pallidum_APC = "BilatPallidum",
    Putamen_APC = "BilatPutamen",
    Thalamus_APC = "BilatThalamus",
    X3rdVentricle_APC = "Bilatv3rdVentricle",
    X4thVentricle_APC = "Bilatv4thVentricle"
  )

  if(!old_name %in%  names(lifebrain_translation)){
    stop(old_name, "not found")
  }

  lifebrain_translation[[old_name]]
}
```

Read in the data and translate.

```
lifebrain <- read_excel("Lifebrain_change_change_n836.xlsx") %>%
  rename(Structure = "...1") %>%
  rename_at(vars(-Structure), ~ map_chr(., translate_lifebrain)) %>%
  mutate(Structure = map_chr(Structure, translate_lifebrain))
## New names:
## * `` -> ...1
```

Perform two-group t-tests in each cluster, testing if the correlation is larger in within than between. Information about the test is printed below.

```
lifebrain_tests <- test_correlations(lifebrain)
## Testing in  Cluster1
##
##  Welch Two Sample t-test
##
## data:  within_correlations and between_correlations
## t = 11.283, df = 15.321, p-value = 3.985e-09
## alternative hypothesis: true difference in means is greater than 0
## 95 percent confidence interval:
##  0.5347269       Inf
## sample estimates:
##  mean of x  mean of y
##  0.4726826 -0.1602484
##
##
##
## Testing in  Cluster2
##
##  Welch Two Sample t-test
##
## data:  within_correlations and between_correlations
## t = 3.7976, df = 50.148, p-value = 0.0001978
## alternative hypothesis: true difference in means is greater than 0
## 95 percent confidence interval:
##  0.1045314       Inf
## sample estimates:
##  mean of x  mean of y
##  0.1363119 -0.0507810
##
##
##
## Testing in  Cluster3
##
##  Welch Two Sample t-test
##
## data:  within_correlations and between_correlations
## t = 2.9104, df = 23.332, p-value = 0.003904
## alternative hypothesis: true difference in means is greater than 0
## 95 percent confidence interval:
##  0.06001752        Inf
## sample estimates:
##    mean of x    mean of y
##  0.139147746 -0.006711137
##
##
##
## Cluster4 has only a single member. No test performed.
##
## Cluster5 has only a single member. No test performed.
```

### LCBC

This correlation matrix is sorted after the order of regions appearing in the figure above. The order is as follows.

```
(lcbc_names <- unname(unlist(clusters)))
##  [1] "Bilatv3rdVentricle"    "Bilatv4thVentricle"    "BilatLatVentricle"
##  [4] "BilatInfLatVentricle"  "BilatBrainStem"        "BilatCerebellumWM"
##  [7] "BilatCerebellumCortex" "BilatThalamus"         "BilatHippocampus"
## [10] "BilatCerebralWM"       "BilatCerebralCortex"   "BilatPutamen"
## [13] "BilatAmygdala"         "BilatAccumbens"        "BilatCaudate"
## [16] "BilatPallidum"
```

Read in the data.

```
lcbc <- read_delim("corrmat_adults_sorted.txt", delim = ",",
                   col_names = lcbc_names) %>%
  mutate(Structure = lcbc_names)
## Parsed with column specification:
## cols(
##   Bilatv3rdVentricle = col_double(),
##   Bilatv4thVentricle = col_double(),
##   BilatLatVentricle = col_double(),
##   BilatInfLatVentricle = col_double(),
##   BilatBrainStem = col_double(),
##   BilatCerebellumWM = col_double(),
##   BilatCerebellumCortex = col_double(),
##   BilatThalamus = col_double(),
##   BilatHippocampus = col_double(),
##   BilatCerebralWM = col_double(),
##   BilatCerebralCortex = col_double(),
##   BilatPutamen = col_double(),
##   BilatAmygdala = col_double(),
##   BilatAccumbens = col_double(),
##   BilatCaudate = col_double(),
##   BilatPallidum = col_double()
## )
```

Perform two-group t-tests in each cluster, testing if the correlation is larger in within than between. Information about the test is printed below.

```
lcbc_tests <- test_correlations(lcbc)
## Testing in  Cluster1
##
##  Welch Two Sample t-test
##
## data:  within_correlations and between_correlations
## t = 17.141, df = 33.991, p-value < 2.2e-16
## alternative hypothesis: true difference in means is greater than 0
## 95 percent confidence interval:
##  0.5812216       Inf
## sample estimates:
##  mean of x  mean of y
##  0.4456900 -0.1991452
##
##
##
## Testing in  Cluster2
##
##  Welch Two Sample t-test
##
## data:  within_correlations and between_correlations
## t = 6.2174, df = 71.134, p-value = 1.534e-08
## alternative hypothesis: true difference in means is greater than 0
## 95 percent confidence interval:
##  0.2575573       Inf
## sample estimates:
##   mean of x   mean of y
##  0.29512893 -0.05674885
##
##
##
## Testing in  Cluster3
##
##  Welch Two Sample t-test
##
## data:  within_correlations and between_correlations
## t = 6.1225, df = 50.806, p-value = 6.677e-08
## alternative hypothesis: true difference in means is greater than 0
## 95 percent confidence interval:
##  0.166707      Inf
## sample estimates:
##  mean of x  mean of y
## 0.28430333 0.05479108
##
##
##
## Cluster4 has only a single member. No test performed.
##
## Cluster5 has only a single member. No test performed.
```

### Summary

The table below summarizes the p-values for each sample.

```
bind_rows(
  UKB = ukb_tests,
  VETSA = vetsa_tests,
  Lifebrain = lifebrain_tests,
  LCBC = lcbc_tests,
  .id = "Dataset"
) %>%
  knitr::kable()
```

| Dataset | Cluster1 | Cluster2 | Cluster3 | Cluster4 | Cluster5 |
| --- | --- | --- | --- | --- | --- |
| UKB | 0 | 0.0000172 | 0.0000006 | NA | NA |
| VETSA | 0 | 0.0000003 | 0.1711224 | NA | NA |
| Lifebrain | 0 | 0.0001978 | 0.0039035 | NA | NA |
| LCBC | 0 | 0.0000000 | 0.0000001 | NA | NA |
